# Structure-guided engineering of type I-F CASTs for targeted gene insertion in human cells

**DOI:** 10.1101/2024.09.19.613948

**Authors:** George D. Lampe, Ashley R. Liang, Dennis J. Zhang, Israel S. Fernández, Samuel H. Sternberg

## Abstract

Conventional genome editing tools rely on DNA double-strand breaks (DSBs) and host recombination proteins to achieve large insertions, resulting in a heterogeneous mixture of undesirable editing outcomes. We recently leveraged a type I-F CRISPR-associated transposase (CAST) from the *Pseudoalteromonas* Tn*7016* transposon (*Pse*CAST) for DSB-free, RNA-guided DNA integration in human cells, taking advantage of its programmability and large payload capacity. *Pse*CAST is the only characterized CAST system that has achieved human genomic DNA insertions, but multiple lines of evidence suggest that DNA binding may be a critical bottleneck that limits high-efficiency activity. Here we report structural determinants of target DNA recognition by the *Pse*CAST QCascade complex using single-particle cryogenic electron microscopy (cryoEM), which revealed novel subtype-specific interactions and RNA-DNA heteroduplex features. By combining our structural data with target DNA library screens and rationally engineered protein mutations, we uncovered CAST variants that exhibit increased integration efficiency and modified PAM stringency. Structure predictions of key interfaces in the transpososome holoenzyme also revealed opportunities for the design of hybrid CASTs, which we leveraged to build chimeric systems that combine high-activity DNA binding and DNA integration modules. Collectively, our work provides unique structural insights into type I-F CAST systems while showcasing multiple diverse strategies to investigate and engineer new RNA-guided transposase architectures for human genome editing applications.

## INTRODUCTION

Canonical CRISPR-Cas systems that have been leveraged for programmable gene editing, such as Cas9 nucleases, cause targeted DNA double-strand breaks (DSBs) that provoke the cell to activate DNA repair mechanisms^1,2^. Non-homologous end joining (NHEJ) is the most efficient repair pathway in human cells, which leads to indel mutations, and although homology-directed repair (HDR) offers the ability to generate precise modifications or insertions, it is inefficient in most cell types, inaccessible in non-dividing cells, and requires large homology arms for each new insertion site^3,4^. Furthermore, HDR efficiencies decrease drastically with insertion size, and aberrant editing pathways that occur at non-negligible frequencies can cause large chromosomal truncations and/or rearrangements^5–10^. Second generation editors, including base and prime editors, employ nickase-variant Cas proteins to bypass DSB intermediates, but indel byproducts still arise and edits are generally restricted to single-base pair (bp) changes or small insertions (<50 bp)^11–14^, thus failing to address the need for large DNA insertion technology. CRISPR-associated transposases (CASTs), on the other hand, leverage a CRISPR-associated DNA targeting module and a transposase effector module that allow for highly specific and programmable insertions, which are both DSB-free and multi-kilobases in size^15–17^.

To date, four CAST subtypes have been characterized in bacteria: type I-B, I-D, I-F, and V-K^15,16,18,19^. These subtypes encode unique architectures for both the targeting and integration steps of the transposition pathway: type I CASTs rely on TnsABC proteins for integration and a multi-subunit complex for DNA targeting that includes TniQ and Cascade components (TniQ-Cascade, hereafter simply QCascade), with Cascade itself comprising 3-5 unique protein components in varying oligomeric states^20–22^; whereas type V-K CASTs rely on only TnsBC for integration^16,23,24^ and a simpler Cas12k-TniQ-S15 co-complex for DNA targeting^25^. Individual homologs within each of these CAST subtypes also vary in sequence identity^26,27^, subunit composition and fusion connectivity^18,24,28^, DNA targeting modules, crRNA guide sequence^18,26,29^, and host factor requirements^17,25,30^, thus representing a diverse pool of potential starting points for tool development. Although type V-K CASTs are more compact systems in terms of coding size, they exhibit multiple undesirable biochemical properties — including reduced specificity^31–33^, low overall editing efficiencies^16,31^, and poor product purity^24,34,35^ — that would necessitate extensive optimization for potential research and therapeutic genome engineering applications. In contrast, type I-F CASTs exhibit highly specific and homogeneous integration products, with demonstrably greater efficiencies than types I-B, I-D, and V-K^15–19,24^.

CAST systems have been the focus of extensive structural efforts using cryogenic electron microscopy (cryoEM) in recent years. The type V-K ShCAST system from *Scytonema hoffmannii* has been systematically investigated^25,36–39^, with a recent report of the holo transpososome architecture that revealed intricacies of the megadalton complex containing Cas12k, TniQ, TnsB, TnsC, single-guide RNA, partial donor and target DNA substrates, and the bacterial host factor S15^39^. Structural studies of type I-B and I-F CASTs have largely focused on the QCascade DNA targeting module and the accessory TnsC ATPase^20,21,40–43^, with no structures of the endonuclease-transposase TnsAB module described to date. Intriguingly, QCascade structures exhibit distinct conformations across different systems: type I-B CASTs feature a single TniQ monomer that recruits TnsC to the Cascade-bound target DNA^21^, whereas type I-F CASTs feature a TniQ homodimer that is stably associated with Cascade^20^. Thus far, two I-F CAST systems from subtypes I-F3a and I-F3b have been deeply characterized — *Vch*CAST (Tn*6677*) and *Asa*CAST (Tn*6900*), respectively — both of which are only distantly related to *Pse*CAST (Tn*7016*), a system that we recently exploited for targeted DNA integration in human cells^17^.

The *Pse*CAST RNA-guided transposase was identified as a lead candidate for human genome engineering applications through a systematic screen of diverse type I-F CAST systems^17^ (**Fig. 1a**). Although our first study reported editing activities that reached single-digit efficiencies at genomic target sites, representing a ∼100-fold improvement over our original candidate, *Vch*CAST, these efficiencies remain limiting for downstream applications. We hypothesized that identifying bottlenecks in the system would inform more targeted rationally engineering, developed several assays to investigate intermediate events and overall integration efficiencies in human cells^17^, and then applied these assays to *Vch*CAST and *Pse*CAST, the only type I-F CASTs shown to successfully perform RNA-guided integration in human cells. Intriguingly, while *Pse*CAST promoted comparatively robust DNA integration, it exhibited markedly weaker DNA binding activity relative to *Vch*CAST. We therefore hypothesized that, alongside parallel efforts to engineer and evolve hyperactive transposase variants, the *Pse*CAST QCascade module would represent a promising focus area to improve DNA targeting and thus editing efficiencies.

**Figure 1.**
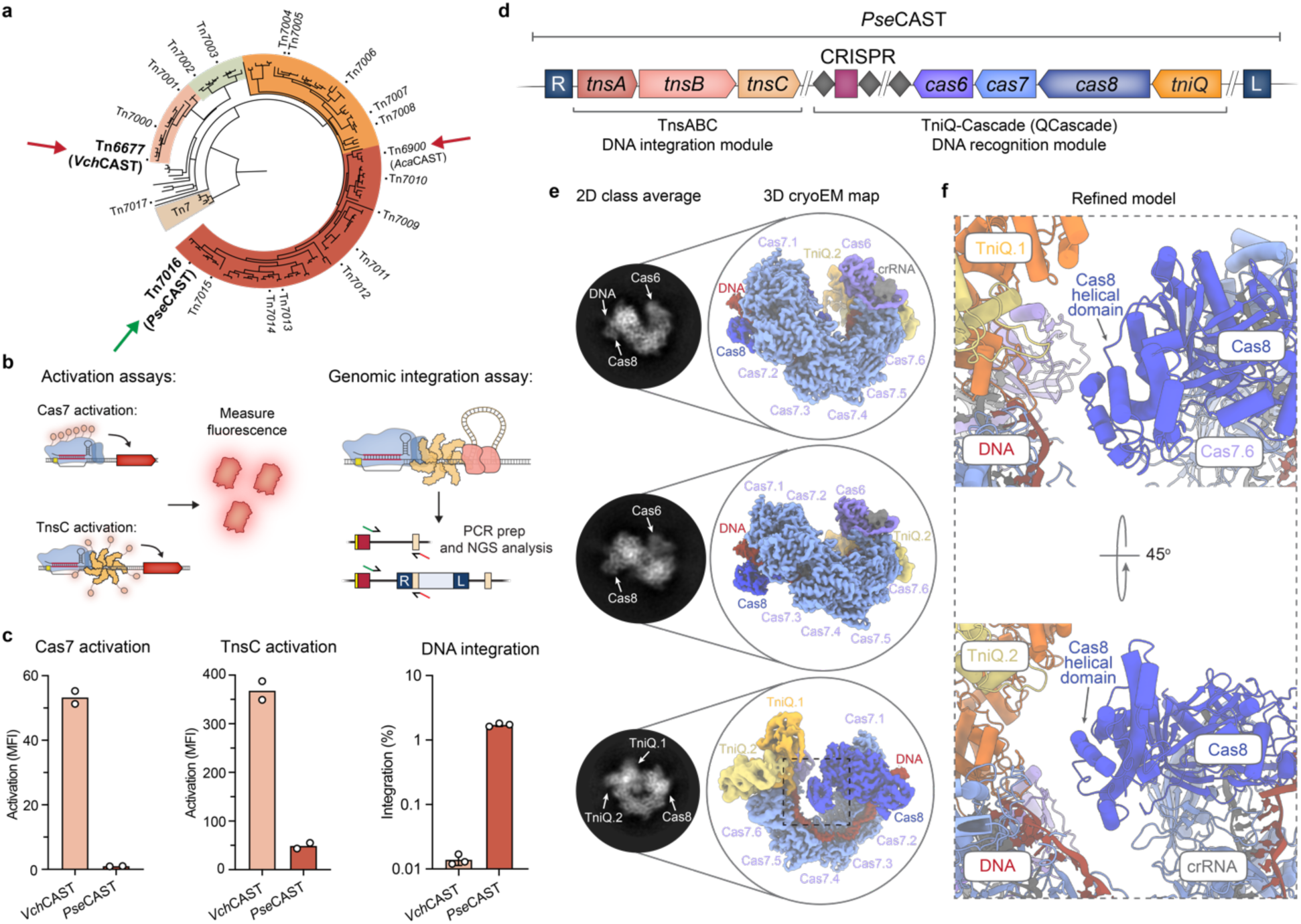
CryoEM structure of the TniQ-Cascade (QCascade) complex from *Pse*CAST. **a,** Phylogenetic tree of type I-F CRISPR-associated transposons (CASTs), adapted from a previous publication^26^. Systems with previously solved QCascade structures are marked with red arrows, while *Pse*CAST is marked with a green arrow. Phylogenetic clades are colored. **b,** Experimental design to investigate both DNA binding and overall integration activities for CAST systems in human cells^17^. DNA binding is extrapolated from two different transcriptional activation assays, one in which VP64 is fused to Cas7 (left), and one in which VP64 is fused to TnsC (right). Overall integration efficiencies are measured via amplicon sequencing. **c,** Comparison of *Vch*CAST and *Pse*CAST across different assays in human cells. Although *Pse*CAST exhibits consistently weak transcriptional activation compared to *Vch*CAST, its absolute integration activity is approximately two orders of magnitude greater. DNA integration data is adapted from a previous publication^17^. **d,** Operonic architecture of *Pse*CAST components from the *Pse*CAST transposon, with genes encoding the QCascade complex labeled accordingly. **e,** Left, dominant reference-free 2D cryoEM class averages. Right, cryoEM densities with colored map regions corresponding to Cas8 (blue), Cas7 monomers 1-6 (light blue), Cas6 (purple), TniQ monomers 1-2 (orange, yellow), crRNA (gray), and target DNA (red) indicated. **f,** Refined model for the Cas8 ɑ-helical domain and its positioning relative to the TniQ dimer interface.

Towards that goal, here we report the cryoEM structure of *Pse*CAST QCascade and the effect of targeted mutations in the PAM- and crRNA-interacting regions on DNA integration. Separately, we leveraged AlphaFold-Multimer to predict protein-protein interactions within the TnsABC co-complex, inspiring the rational design of novel chimeric CAST systems that enable divergent DNA targeting and DNA integration modules to be combined into a single functional system. Collectively, this work establishes multiple biochemically- and structurally-guided approaches to engineer CAST systems for improved editing efficiencies in human cells.

## RESULTS

### CryoEM structure of *Pse*CAST QCascade complex

We previously demonstrated that *Vch*CAST and *Pse*CAST, two distantly related type I-F CASTs^17,26^, exhibit distinct DNA binding and integration efficiencies (**Fig. 1a-c**). Given our previous mechanistic and structural studies of the QCascade complex from *Vch*CAST^20,40^, we hypothesized that structure-guided engineering of the *Pse*CAST QCascade complex might reveal novel interactions and open a path to improve overall integration efficiencies. We therefore purified recombinant *Pse*QCascade after carefully optimizing the expression vector design (**Supplementary Fig. 1**) and set out to determine the cryoEM structure.

We incubated the purified *Pse*QCascade complex, which is expected to comprise a 1:6:1:2:1 stoichiometry of Cas8:Cas7:Cas6:TniQ:crRNA components (**Fig. 1d**), with a double-stranded DNA (dsDNA) substrate containing a 32-bp target sequence and 5′-CC-3′ PAM, and then subjected the sample to electron microscopy. Preliminary cryoEM experiments revealed a homogeneous behavior with multiple views and no apparent disassembly (**Supplementary Fig. 2a**), and the overall architecture was consistent with other type I-F QCascade complexes, comprising six Cas7 monomers (named hereafter Cas7.1 to Cas7.6) that form a pseudo-helical assembly coating the crRNA molecule (**Fig. 1e**). The Cas8 protein contains two domains: a bulky domain that interacts with Cas7.1 and binds the crRNA 5′ end and PAM sequence, and a second ɑ-helical domain that exhibited a dynamic behavior (**Fig. 1f**). Towards the crRNA 3′ end (hereafter PAM-distal region), the RNA hairpin is stabilized by Cas6, which also binds the TniQ dimer. Preliminary maps exhibited greater mobility for the TniQ dimer compared to other QCascade components (**Supplementary Fig. 2b,c**). The quality of the maps approaching the TniQ dimer region degrades rapidly, contrasting the excellent map quality for the PAM-adjacent region (**Supplementary Fig. 2d**). Multibody approaches in Relion4 improved the overall resolution, with approximately 2.6 Å and 3.0 Å resolution estimates in the PAM-proximal and PAM-distal regions, respectively (**Methods**).

To further characterize the dynamics of the system and confirm the existence of novel interactions, we complemented our multibody analysis in Relion4 with cryoDRGN^44^, a machine-learning approach for cryoEM analysis (**Supplementary Fig. 3**). CryoDRGN revealed multiple populations of the complex, with the TniQ dimer populating a wide range of positions relative to the rest of the complex that pivot around Cas6 and Cas7.6. The dimer adopts an ‘open’ conformation that lacks any direct interactions with Cas8, as well as multiple intermediate, ‘closed’ conformations that approach the tip of the Cas8 ɑ-helical domain (**Supplementary Fig. 3b**). In a recent structure of a homologous QCascade complex bound to target DNA, the Cas8 ɑ-helical domain exhibits a different conformation, almost perpendicular to the inner face of the TniQ dimer and aligned with the bulky domain of Cas8^22^. In our dataset, we did not observe this extended conformation and instead detected alternative TniQ-Cas8 interactions that are established between the most distal end of the TniQ dimer and the apical part of the Cas8 ɑ-helical domain, which were revealed through low-pass filtered maps (**Supplementary Fig. 3c**). Both the TniQ dimer and the Cas8 ɑ-helical domains remain in parallel configurations, with only marginal contacts at the periphery of the complex. Despite the apparent flexibility in this interaction (**Supplementary movie 1 and 2**), the Cas8 ɑ-helical domain is essential for RNA-guided DNA integration activity, as revealed by the complete loss of human cell activity when we replaced the domain with a flexible glycine-serine linker (**Supplementary Fig. 4**).

### Stabilizing protein-RNA and protein-protein interactions

The overall architecture of the TniQ dimer is similar to the *Vch*CAST QCascade dimer^20^, with an antiparallel head-to-tail configuration, forming a compact unit that laterally approaches the interface formed by Cas6 and Cas7.6 (**Fig. 2a**). The C-terminal domain of one TniQ monomer interacts with Cas6, and the N-terminal domain of the other TniQ monomer interacts with Cas7.6. At the core of this four-fold interface, the crRNA appears to play a critical role, with residues 40– 45 establishing multiple RNA-protein stacking interactions (**Fig. 2b,c**).

**Figure 2.**
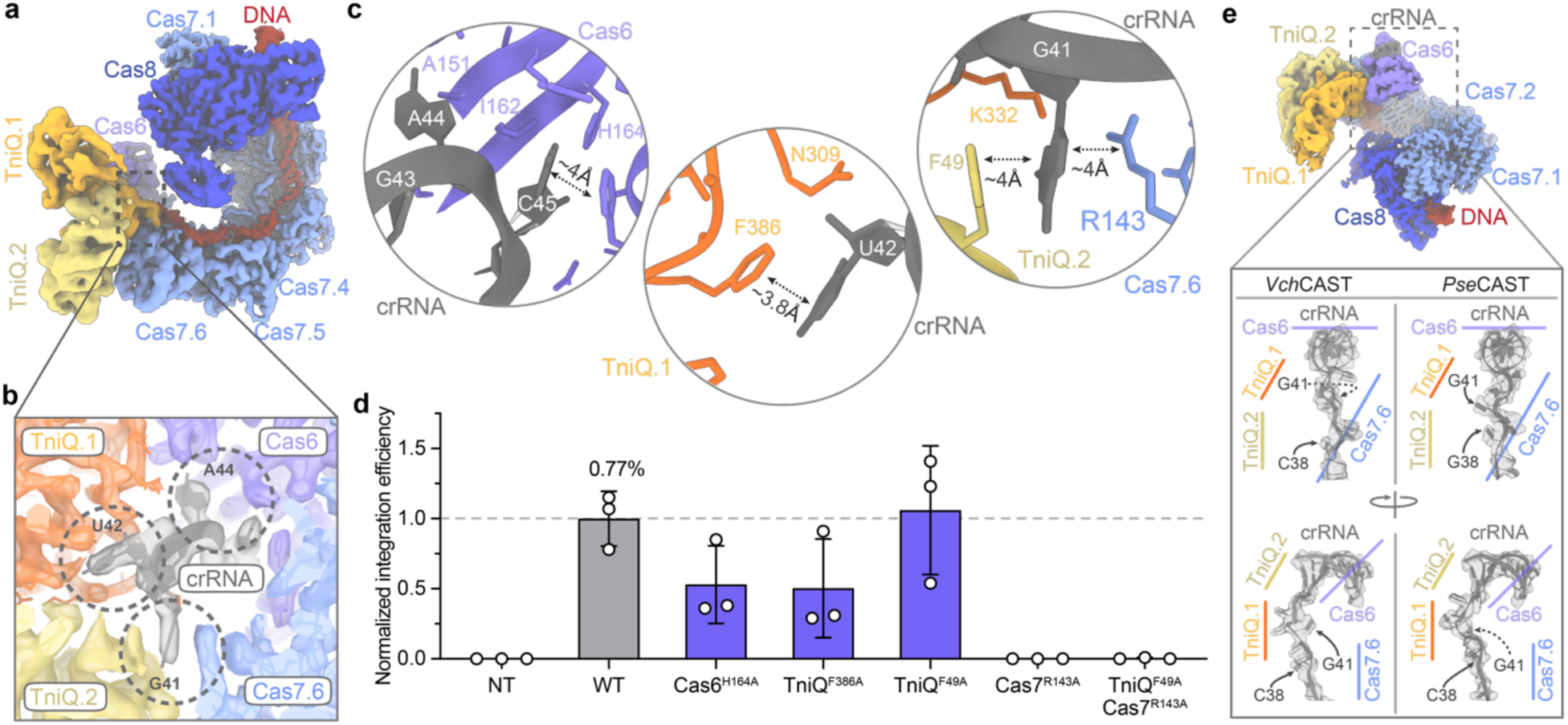
The role of crRNA in the PAM-distal region of *Pse*QCascade. **a,** Overall view of the cryoEM reconstruction of the *Pse*CAST QCascade complex. **b,** Magnified view of the dashed region in **a**, highlighting the cryoEM density (colored and semi-transparent) for interactions between the indicated crRNA nucleotides and protein subunits. **c,** Magnified view of the dashed regions in **b**, highlighting interactions between the crRNA and Cas6 (left), TniQ.1 (middle), and both TniQ.2 and Cas7.6 (right). Key interacting residues are labeled. **d,** Normalized RNA-guided DNA integration efficiency at *AAVS1* in HEK293T cells, as measured by amplicon sequencing. The indicated alanine mutations were designed to perturb specific RNA-protein interactions highlighted in **c**, and were compared to WT. NT, non-targeting crRNA. Data are shown as mean ± s.d. for n=3 biologically independent samples. **e,** Comparison of the crRNA conformation within the PAM-distal region, adjacent to the site of RNA hairpin stabilization by Cas6, for *Vch*CAST (PDB: 6PIJ) and *Pse*CAST (this study). The region around nucleotide G41 exhibits a distinct configuration for *Pse*CAST, likely affecting the behavior of the adjacent TniQ dimer.

We hypothesized that crRNA interactions with Cas6, Cas7.1, TniQ.1, and TniQ.2 are crucial for robust QCascade complex formation, and that disrupting them would prevent transposase recruitment and abolish integration activity. We therefore introduced alanine point mutations to disrupt nucleobase-side chain stacking interactions and investigated the resulting effects in human genomic DNA integration assays. Alanine substitutions to Cas6 and TniQ residues contacting the crRNA were well tolerated, whereas a Cas7 R143A mutation (Cas7^R143A^) abolished integration activity (**Fig. 2d**). The crRNA trajectory in the hinge region between Cas7.6 and Cas6 differs in *Pse*CAST and *Vch*CAST (**Fig. 2e**), and *Pse*CAST crRNA residue G41 seems to play a key role as an interaction “hub,” establishing coincident contact with TniQ.1, TniQ.2, and Cas7.6 by adopting a unique, extruded conformation.

We next explored protein-protein interactions that we similarly hypothesized would contribute to QCascade function, in part by playing a role in downstream transposase recruitment to the target site. The first of these interactions involved a hydrophobic patch on Cas6 cradling hydrophobic residues in the loop connecting TniQ.1 α-helices W262–K275 and F312–S327 (**Fig. 3a,b**), which is conserved across homologous QCascade complexes, with minor variations. Specifically, a hydrophobic residue in the TniQ.1 connecting loop (I282 in *Pse*CAST, V270 in *Vch*CAST) inserts deeply into the Cas6 hydrophobic patch to anchor the TniQ monomer to the Cascade module (**Fig. 3c**). The cradle structure of this interaction potentially acts as a pivot point, facilitating dynamic TniQ movement. Disruption of these hydrophobic interactions via introduction of charged arginine residues in either TniQ or Cas6 led to a marked reduction in integration efficiencies (**Fig. 3d**). The other TniQ monomer (TniQ.2) interacts electrostatically with Cas7.6 via α-helix Y33–L47 and adjacent residues (**Fig. 3e**). Given the multimeric assembly of Cas7 monomers along the crRNA, loop regions observed to interact with TniQ.2 may have pleiotropic functions, possibly participating in Cas7 monomer-monomer interactions (**Supplementary Fig. 5**). With the goal of selectively perturbing Cas7.6-TniQ.2 interactions to investigate its importance, we avoided mutagenizing residues that might affect the Cas7 monomer-monomer contacts and thus focused on loops A and B (**Supplementary Fig. 5b**). Alanine mutations within the TniQ-interacting regions abolished DNA integration, whereas several mutations within Cas7 had surprisingly little to no impact on overall DNA integration activity (**Fig. 3f**).

**Figure 3.**
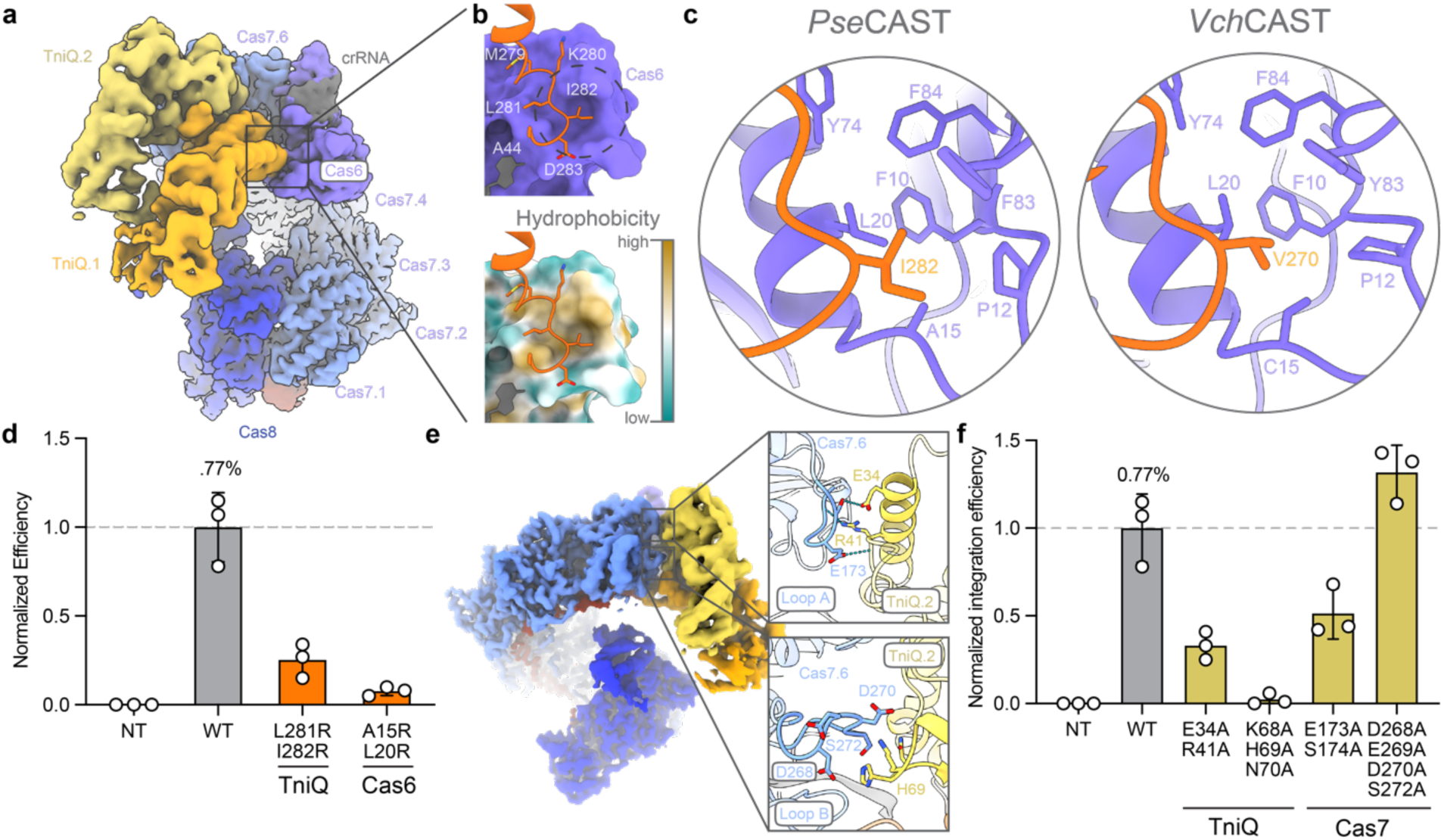
TniQ recruitment to the Cas6-Cas7.6 interface of Cascade requires hydrophobic and electrostatic interactions. **a,** Overall view of the *Pse*CAST QCascade complex, oriented to highlight the TniQ dimer (dark/light orange). **b,** Magnified view of the region indicated in **a**, showing how TniQ.1 (dark orange) interacts with a hydrophobic cavity on Cas6. The two visual renderings are colored either by Cas6 surface (purple, top) or hydrophobicity (bottom). **c,** Comparison of the hydrophobic interactions between TniQ.1 and Cas6 in *Pse*CAST (left) and *Vch*CAST (right, PDB: 6PIJ), with residues labeled. **d,** Normalized RNA-guided DNA integration efficiency at *AAVS1* in HEK293T cells, as measured by amplicon sequencing. The indicated arginine point mutations were designed to perturb TniQ.1-Cas6 hydrophobic interactions. NT, non-targeting crRNA. **e,** Magnified views of hydrogen bonding (top) and electrostatic (bottom) interactions between Cas7.6 (blue) and TniQ.2 helix (yellow). **f,** Normalized RNA-guided DNA integration efficiency at *AAVS1* in HEK293T cells, as measured by amplicon sequencing. Alanine mutations perturbing Cas7.6-TniQ interactions are generally tolerated. Data in **d, f** are shown as mean ± s.d. for n=3 biologically independent samples.

### Protein engineering modulates PAM stringency and improves DNA integration

In comparison to other type I-F CASTs, *Pse*CAST exhibits a remarkably flexible PAM preference, with almost no sequence preference at both the -1 and -2 positions in *E. coli* transposition assays^26^; this property may lead to a dramatic increase in the effective search space for the 32-bp guide. Inspired by previous work investigating CRISPR-Cas9 activity and PAM search space^45^, we hypothesized that inefficient DNA targeting due to a flexible PAM preference may represent a rate-limiting step in RNA-guided DNA integration, especially within the cellular milieu of human cells, whose genome is ∼1000× larger than *E. coli*. We therefore set out to specifically engineer QCascade variants that might exhibit altered PAM specificity and thus direct altered DNA integration efficiencies.

After leveraging the excellent quality of our cryoEM map in the area surrounding Cas8, we identified two hydrophobic alanine residues at the center of the PAM-interacting region. In contrast, systems with stricter PAM preferences — *Vch*CAST, *Asa*CAST, and *Pae*Cascade from a *Pseudomonas aeruginosa* type I-F1 CRISPR-Cas system^26,46^ — feature polar residues at the equivalent positions, which allow for hydrogen bonding with specific PAM nucleotides (**Fig. 4a,b, Supplementary Fig. 6a**). Based on these observations, we reasoned that mutation of A143 and A144 to residues with greater hydrogen bonding potential might improve PAM stringency, reduce the effective search space, and result in more efficient DNA targeting. We also chose to mutagenize residues 125–127, as this region also interacts with the PAM (**Fig. 4b, Supplementary Fig. 6a**). We analyzed the sequence conservation at these PAM-interacting regions and compared *Pse*CAST to other Cascade homologs that have previously exhibited either robust DNA integration activity or stringent PAM preferences (**Supplementary Fig. 6b,c**). Collectively, we designed fifteen Cas8 variants with PAM-interacting mutations, varying from single point mutations at A243 or A244 to larger mutations in which the entire PAM-interacting region was grafted from a homolog.

**Figure 4.**
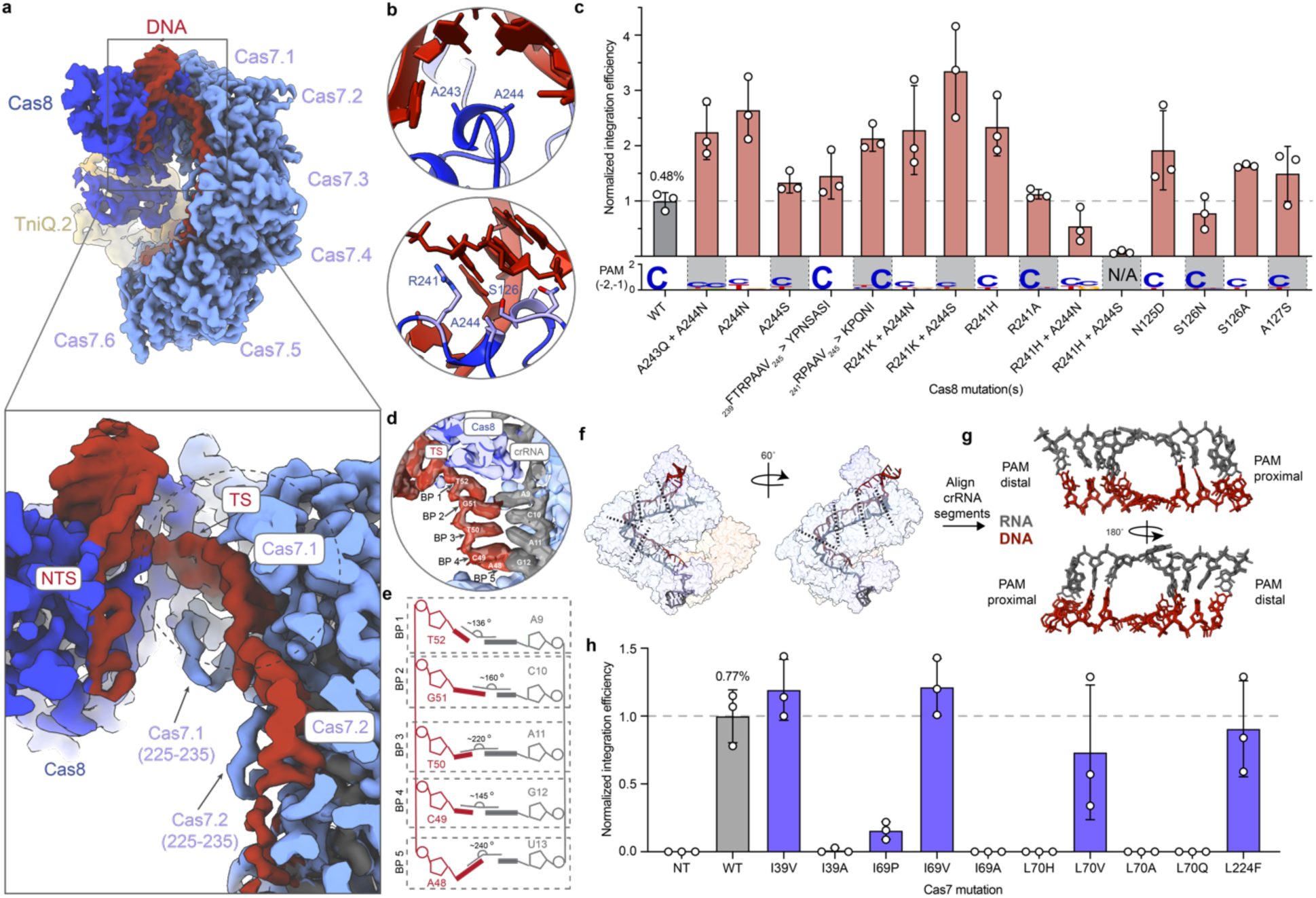
Structural and functional consequences of PAM and target DNA recognition by *Pse*QCascade. **a, Top,** overall view of the *Pse*CAST QCascade complex, oriented to highlight the target DNA recognition. **Bottom,** magnified views of the PAM binding pocket, with Cas8 and DNA shown in blue and red, respectively. Residues A243 and A244 lack any base-specific, hydrogen-bonding interactions with the DNA. **b,** Normalized genomic integration efficiencies at AAVS1 for the indicated Cas8 mutants (top), plotted above the WebLogo for PAM preferences in the -1 and -2 positions (bottom) derived from integration into pTarget. (For additional PAM specificity data, see **Supplementary Fig. 6e.**) Integration efficiency data are shown as mean ± s.d. for n=3 biologically independent samples. **c,** Magnified view of the experimental cryoEM density map around Cas7.1 and Cas7.2, showing interactions with the crRNA (gray) and DNA target strand (TS, red). NTS, DNA non-target strand. **d,** Overlay of the refined atomic model and cryoEM density (semi-transparent) for the seed region of QCascade bound to the DNA target strand. **e,** Schematic representation showing angles for the first five RNA-DNA base pairs (BP 1– 5) within the R-loop. **f,** View of the RNA-DNA heterduplex at right, highlighting the unfavorable base-pairing surrounding flipped out nucleobases within the first 18 base pairs of the R-loop. **g,** Magnified view of the RNA-DNA heteroduplex segments aligned at the flipped out base pair, revealing consistent unfavorable angles at the adjacent base pairs. **h,** Normalized RNA-guided DNA integration efficiency at *AAVS1* in HEK293T cells for the indicated Cas7 mutations, as measured by amplicon sequencing. Data are shown as mean ± s.d. for n=3 biologically independent samples.

We quantified changes in PAM preference by performing an episomal PAM library screen in HEK293T cells, in which a target plasmid (pTarget) contained an *AAVS1* target site directly downstream of a randomized 4-bp PAM library (**Supplementary Fig. 6d**). After transiently transfecting cells with pTarget, pDonor, and all the necessary protein-RNA expression vectors, we isolated plasmid DNA, sequenced the PAM motifs from all successful integration products, and constructed a consensus motif for each Cas8 variant; in parallel, we also quantified absolute integration efficiencies at the genomic AAVS1 site, which contains a 5′-CC-3′ PAM (**Fig. 4c**). The results revealed that certain mutations led to improvements in integration efficiencies by as much as 3.5-fold, but without a clear correlation between PAM stringency and overall genomic integration activity (**Fig. 4c**). For example, the variant with the greatest improvement in integration activity, Cas8^R241K,A244S^, actually exhibited a reduced PAM preference, compared to the stronger preference for cytidine in the -2 position with WT Cas8 (**Fig. 4c, Supplementary Fig. 6e**). Interestingly, Cas8^A243Q,A244N^ exhibited decreased PAM preference, whereas when we grafted the entire PAM region from a type I-F1 system (_241_RPAAV_245_>KPQNI), the resulting mutant restored a strong preference for cytidine at the -1. Mutations within the upstream PAM-interacting region (residues 125-127) showed moderate improvements on integration activity, with either unchanged or moderately reduced PAM stringency (**Fig. 4c**). A Cas8^R241A^ mutant with disrupted ‘R-wedge,’ which normally forms stacking interactions with the -1 PAM position to help unwind dsDNA^47,48^, unexpectedly exhibited both WT integration efficiencies and PAM stringency (**Fig. 4c**).

Together, mutational profiling of the PAM-interacting region revealed key residues whose mutation improved integration efficiencies, but the combination of PAM specificity and integration activity results failed to support the hypothesis that PAM promiscuity is a key bottleneck towards achieving higher efficiency *Pse*CAST integration activity in human cells (**Fig. 4c, Supplementary Fig. 6e**).

We also focused on PAM-proximal interactions with the upstream double-stranded DNA region as another potential point of engineering and optimization. Previous work on canonical type I-F1 defense systems revealed key interactions between dsDNA and the N-terminal region of Cas8^47–49^, with a positively charged vise domain undergoing a conformational change to ‘clamp’ onto the PAM-adjacent sequence in a non-specific fashion. When comparing *Pse*Cas8 (from type I-F3 *Pse*CAST) to *Pae*Cas8 (from type I-F1 *Pae*Cascade; **Supplementary Fig. 7a**), we observed a markedly different conformation of the N-terminus, with the vise domain absent. Given this potential deficiency, we hypothesized that substituting the *Pae*Cas8 vise domain in *Pse*Cas8 could improve DNA binding affinity and thus CAST activity. However, a thorough screening of chimeric Cas8 constructs for human cell integration activity revealed a clear intolerance of *Pse*Cas8 to sequence perturbations in this region (**Supplementary Fig. 7b**). We pursued additional synthetic strategies to improve DNA binding of *Pse*QCascade by fusing a variety of DNA-binding domains to the *Pse*Cas8 N-terminus of *Pse*Cas8 (**Supplementary Fig. 7c**), inspired by engineering strategies previously applied to polymerases^50,51^, reverse transcriptases^52^, and ligases^53^. However, these fusions exhibited no improvement relative to WT, and in some cases reduced overall integration efficiencies (**Supplementary Fig. 7c**). Collectively, these experiments suggest that either the DNA binding affinity of *Pse*Cas8 is not a critical bottleneck in the overall transposition pathway, or that the tested variants fail to improve upon the WT activity in this regard.

### Unfavorable nucleobase positioning along the RNA-DNA heteroduplex

Cascade complexes bind the target DNA by forming a discontinuous RNA-DNA heteroduplex in 6-bp segments^47,54^, and we could clearly resolve RNA-DNA base pairs for the first 4 segments engaged by Cas7 monomers within the *Pse*QCascade complex, but the remaining two segments featured weaker RNA density and no DNA density. Density for the RNA-DNA heteroduplex across the first 3 segments (crRNA residues 9 to 26) was exceptionally good, with clear separation within base pairs and features compatible with a local resolution beyond 3 Å. We were therefore able to accurately model RNA-DNA interactions to a high level of confidence in these regions of the map. The resulting view revealed peculiarities in the base-pair geometry, with acute divergence from ideal values in some base pairs. The third and fourth base pair within each segment exhibited severe deviation from ideal planarity values (buckling), while the first and fifth base pair exhibited exacerbated propeller twist deviations. Only the second base pair across distinct segments exhibited geometric and hydrogen-bonding distance values closer to energetically favored conditions (**Fig. 4d-h**).

Type I-F Cascade complexes bind the target DNA, such that the two-stranded β-sheet ‘finger’ motif of each Cas7 monomer engages the crRNA to flip out every sixth nucleotide of the 32-nt spacer, thereby preventing RNA-DNA basepairing^20,47^. We hypothesized that finger motif residues involved in this nucleotide dislocation might promote the consistent distortion of adjacent base pairs, and to explore this effect, we introduced Cas7 mutations intended to relax this distortion, hoping to promote energetically favorable hydrogen-bonding geometries and stabilize the RNA-DNA heteroduplex. Taking advantage of the high local resolution around this region (**Supplementary Fig. 8a,b**), we identified numerous bulky hydrophobic residues —including I69, L70, and L224 — that were not highly conserved across nearby homologs (**Supplementary Fig. 8c**), and subjected them to site-directed mutagenesis.

After generating the desired Cas7 mutations, we performed genomic DNA integration experiments in HEK293T cells at the AAVS1 locus (**Fig. 4i**). Intriguingly, the Cas7 heteroduplex-interacting residues, though not highly conserved, appeared to have low tolerance for mutations. While Cas7^L224F^ and various valine mutations exhibited near-WT integration efficiencies, all other mutations, including Cas7^I69P^, resulted in detrimental impacts on DNA integration (**Fig. 4i**). Intriguingly, L70H, which would theoretically recapitulate a stacking interaction observed in our previous *Vch*CAST structure^20^, completely ablated integration activity (**Fig. 4i**). Together, the intolerance to perturbations in the Cas7 finger domain suggests these nucleobase kinking interactions may in fact be necessary for proper successful DNA integration.

### Structure-based engineering of chimeric CAST systems

Rational engineering of *Pse*QCascade yielded only moderate improvements in integration activity, suggesting a non-trivial path forward to overcome the apparently weak DNA binding activity in human cells^17^. Although recent studies shed light on the kinetics of Cascade target search and recognition^55,56^, the intermediate steps of Cascade complex formation, TniQ-Cascade association, and 3D-diffusion remain poorly understood, particularly in human cells. *Pse*CAST was originally identified through a homolog screen that investigated both overall integration activity and several subunit-specific properties: crRNA processing, TnsB-donor DNA interactions, and QCascade and TnsC-mediated transcriptional activation^17^. Through this screening process, *Vch*CAST (Tn*6677*) and *Pse*CAST (Tn*7016*) were the only two systems that yielded detectable DNA integration in human cells, despite exhibiting distinct subunit-specific activities. Based on these results, we hypothesized that natural CAST systems may be unlikely to possess optimal human cell properties across all recombinant components, and we therefore set out to design chimeric CAST systems that would enable ‘crosstalk’ between otherwise orthogonal components. Our specific goal was to combine the most active DNA targeting and DNA integration machineries derived from divergent CASTs (**Fig. 5a**).

**Figure 5.**
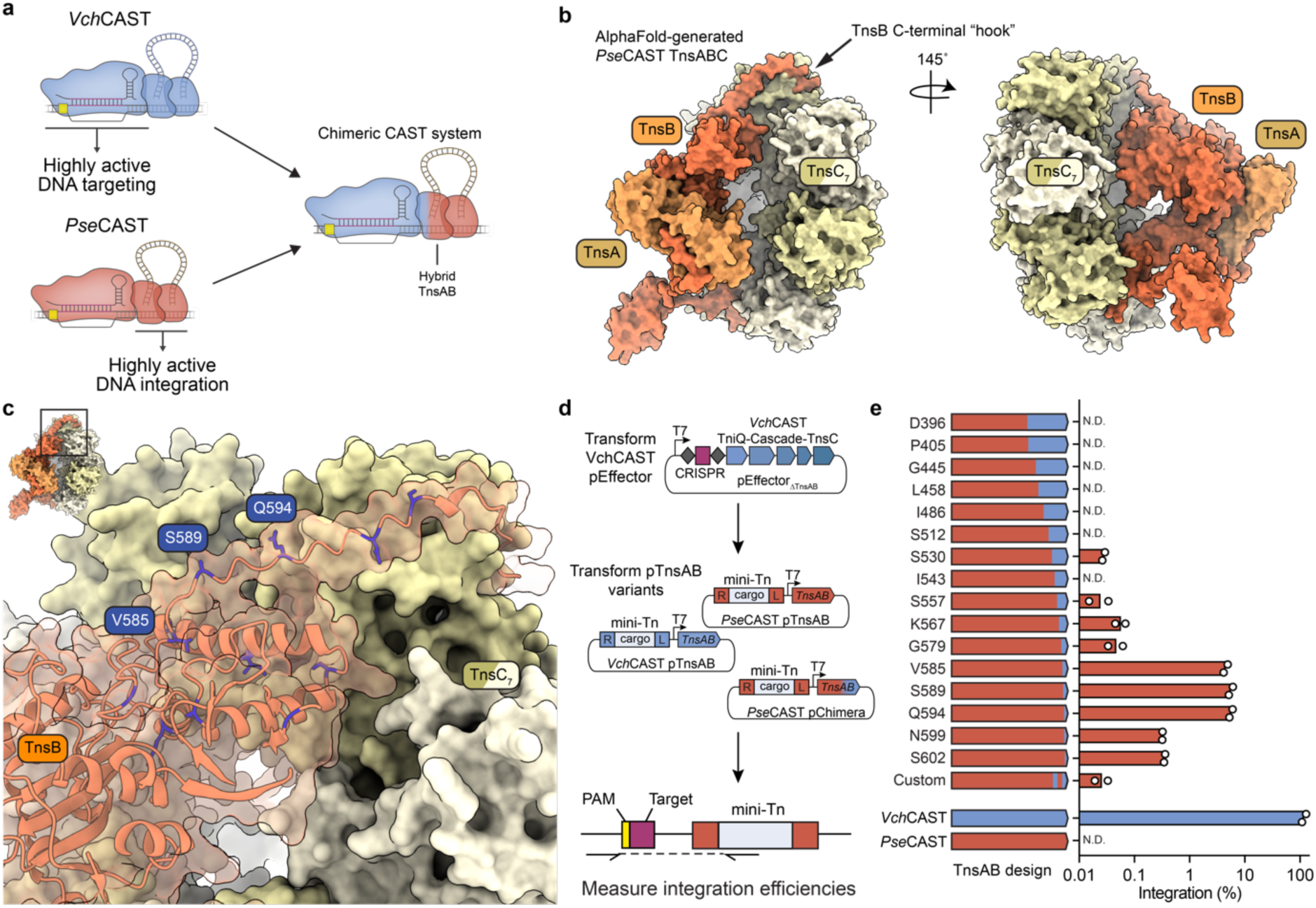
AlphaFold-guided engineering of TnsABC to generate chimeric CAST systems. **a,** Schematic showing the approach to generate a chimeric CAST system by combining optimal DNA targeting and DNA integration machineries from distinct CAST systems. **b,** AlphaFold-generated structure prediction of the TnsABC co-complex from *Pse*CAST. The C-terminal “hook” region of TnsB that putatively interact with TnsC is marked. **c,** Visualization of select TnsB graft points within the predicted *Pse*TnsABC structure. Residues where *Pse-Vch* chimerism was introduced are colored in blue, and the three top performing graft points (V585, S589, Q594; *Pse*TnsB numbering) from panel **e** are labeled. **d,** Experimental workflow to test chimeric TnsAB constructs for RNA-guided DNA integration activity. *E. coli* BL21(DE3) cells containing a pEffector encoding *Vch*QCascade and *Vch*TnsC were transformed with a plasmid encoding a mini-transposon (mini-Tn) and TnsAB, with TnsAB derived from either *Vch*CAST, *Pse*CAST, or a chimeric combination thereof. Integration efficiency was measured by qPCR (bottom). **e,** DNA integration efficiencies for each tested TnsAB chimera. The amino acid listed represents the position at which the reading frame was grafted from *Pse*TnsB (red) to *Vch*TnsB (blue). “Custom” denotes a variant in which multiple different *Vch*TnsB sequences were substituted (see **Supplementary Table 2** for details). Data are shown as mean for n=2 biologically independent samples.

To identify robust DNA targeting homologs, we tested DNA binding activity across 20 type I-F CASTs via transcriptional repression in *E. coli*^40,57^ (**Supplementary Fig. 9a**). Surprisingly, QCascade complexes from only two systems — *Vch*CAST and Tn*7005* — exhibited RFP repression under the tested conditions, with only weak activity from *Pse*CAST and Tn*7000* (**Supplementary Fig. 9b**). Yet when we tested the overall DNA integration activity of *Vch*CAST and *Pse*CAST at the exact same sites used for transcriptional repression, we again observed greater integration activity for *Pse*CAST, mirroring our results in human cells^17^ (**Supplementary Fig. 9c**). This reinforced the conclusion that the weak DNA targeting activity of *Pse*CAST may impose a lower ceiling on achievable DNA integration efficiencies in diverse cell types, despite having co-evolved with a highly active transposition (TnsABC) module.

We sought to address this potential bottleneck by combining the TnsABC machinery from *Pse*CAST with the QCascade machinery from *Vch*CAST. We previously demonstrated that intrinsic CAST modularity precludes simply mixing and matching components from evolutionary diverse systems^26^, but we were emboldened to attempt a more nuanced approach by taking advantage of recent high-resolution structures^21,39^, predicted structures via structural alignments^58^, and AlphaFold-multimer^59^ predicted structures. (**Fig. 5b, Supplementary Fig. 10**). In particular, a model for the putative TnsABC co-complex from *Pse*CAST featured the expected heptameric arrangement of TnsC, similar to our empirical structures for *Vch*CAST^40^, while also revealing predicted interactions between *Pse*TnsC and the C-terminus of *Pse*TnsB that were reminiscent of the TnsB ‘hook’ described for type V-K ShCAST^37,39^ (**Fig. 5b, Supplementary Fig. 10a**). This model, in conjunction with experimentally determined type V-K structures and biochemical studies of Tn*7*^60^, led us to speculate that the C-terminal tail of TnsB functions a key mediator of TnsC interactions, and that the specificity of CAST transpososome assembly would be dictated in part by cognate TnsB-TnsC interactions. Importantly, we hypothesized that reengineering this interaction would enable the TnsAB and donor DNA components from one CAST system to be combined with the QCascade and TnsC components from an orthogonal CAST system.

To test this hypothesis, we designed 16 chimeric TnsAB constructs in which different lengths of the *Pse*TnsB C-terminus were substituted with corresponding residues from the *Vch*TnsB C-terminus (**Fig. 5c**). These variants were then screened for RNA-guided DNA integration activity in *E. coli*, in conjunction with *Vch*QCascade and *Vch*TnsC, but with a pDonor containing transposon ends compatible with *Pse*TnsB (**Fig. 5d**). As expected, given our previous work^26^, WT *Pse*TnsAB, lacking any chimeric substitutions, showed undetectable activity when combined with *Vch*CAST DNA targeting machinery (**Fig. 5e**). Remarkably, however, several chimeric TnsAB designs were able to robustly rescue activity, showing up to ∼10% integration efficiencies (**Fig. 5e**). These designs, which only reprogrammed 20 – 29 amino acids in the C-terminus of *Pse*TnsAB exhibited graft points between the *Pse* and *Vch*TnsB sequence in an unstructured region that links the “hook” region of the C-terminus to the remainder of the protein sequence (**Fig. 5c**); furthermore, when comparing this region to solved type V-K complexes, it is located in a similar region as the 52-residue long “flexible linker” that was unresolved^39^. Together, we conclude that substitutions in this region minimize disruptions to the overall protein fold, while nonetheless providing a chimeric hook that is compatible for cognate interactions with *Vch*TnsC.

We next set out to investigate if this chimeric design is reciprocal; that is, can we rescue DNA integration activity when combining *Pse*QCascade and *Pse*TnsC with a chimeric *Vch*TnsAB design? After designing and cloning similar constructs, we were indeed able to detect integration activity with the converse combination (**Supplementary Fig. 11a**). Furthermore, when we applied these chimeric designs to a broader range of homologous TnsAB variants and their cognate mini-Tn donor substrates, we also observed integration activity for chimeric designs derived from additional transposon variants, denoted Tn*7005* and Tn*7015*^26^. Intriguingly, TnsAB chimeras derived from Tn*7010* and Tn*7011* showed no evidence of activity (**Supplementary Fig. 11b**), suggesting that some CASTs may require targeted screening to identify tolerable chimeric graft points. Next, we explored whether this engineering approach could also generate compatible chimeras between divergent CRISPR-associated transposons, candidate Type I-F (*Vch*CAST) and Type V-K (*Sh*CAST) systems, each of which comprise distinct transposase architectures and likely arose from unique domestication events^23^. TnsB variants derived from *Sh*CAST exhibited low, but detectable levels of activity as well (**Supplementary Fig. 11b,c**); when we investigated the transposon insertion orientation preference for type I/V CAST chimeras, we observed that chimeras in which the TnsB was derived from *Sh*CAST exhibited a “T-LR” insertion preference, as typically observed in previous *Sh*CAST studies^16,35^, while type I-F CASTs exhibit a “T-RL” preference^15,26^ (**Supplementary Fig. 11d**). Together, these results reveal that rational, structure-guided engineering of precise regions of CAST systems can overcome the natural orthogonality of diverse systems, enabling novel genome editing designs.

## DISCUSSION

The unexpected paradox of poor DNA binding and strong overall integration activity of *Pse*CAST (**Fig. 1b,c, Supplementary Fig. 9**), inspired us to determine cryoEM structures of *Pse*QCascade and pursue rational engineering methods to improve DNA targeting. Given the unique phenomenon among CAST systems to harbor ‘homing’ crRNAs that target conserved, often essential, genes within the host genome^18,26,28,29^, CAST-derived CRISPR modules may have been naturally selected for weak DNA binding relative to their defense-associated CRISPR-Cas counterparts, thereby reducing transcriptional repression of these essential genes. This possibility underscores the need to develop a comprehensive understanding of all molecular requirements and intermediate steps within the CAST transposition pathway.

The structure of *Pse*QCascade resembles previously determined DNA-bound type I-F CAST structures^20,22^, but several knowledge gaps still limit a complete understanding of the mechanistic requirements for RNA-guided transposition. First, the functional relevance of the Cas8 helical bundle remains a mystery. When comparing between three distinct, DNA-bound QCascade structures^20,22^, three different conformational states of the helical bundle have been observed: a state in which the domain is unresolved, suggesting a conformationally dynamic mode related to the open versus closed state of the overall QCascade complex^20^; a state in which the domain is resolved, with close contact to the PAM-distal DNA^22^; and a state in which the helical bundle is resolved but does not contact TniQ or the PAM-distal DNA (**Fig. 1e,f**). Despite this heterogeneity, our deletions experiments clearly indicate that the helical bundle is crucial for overall DNA integration to occur (**Supplementary Fig. 4**). Another area that will require future study is the manner in which the QCascade complex binds TnsC, since these interactions have not yet been captured for a type I-F CAST system; mutations in Cas7 that theoretically disrupt Cas7.6 interactions with TniQ.2 appear to be tolerated (**Fig. 3e,f**). Although unexpected, this lends credence to the possibility that only one of the two TniQ monomers present in type I-F CAST complexes interacts with TnsC, which is supported by similar CAST structures from type I-B and type V-K systems in which only one TniQ is present with TnsC at the target site (**Supplementary Fig. 10**)^21,25,39^. Further *in vitro* biochemical studies, combined with structural insights into the holo transpososome, will be necessary to shed light on these mechanistic aspects, including the extent to which the helical bundle may regulate TnsC recruitment, and thus the targeting discrimination between on- and off-target sites during CAST transposition^40^.

Beyond defining structural requirements for transposition, our QCascade structure revealed potential targets for rational engineering, most notably within the PAM interacting regions of Cas8. The presence of alanine residues at this interface, rather than polar residues, differentiates *Pse*CAST from homologous type I-F CAST systems (**Supplementary Fig. 6a**). Interestingly, one of these homologous systems — *Vch*CAST — exhibited higher DNA binding activity than *Pse*CAST in both human cells and *E. coli* (**Fig. 1c, Supplementary Fig. 9**), leading us to hypothesize that reinstating polar residues might stabilize DNA-protein interactions, thereby increasing DNA binding activity and integration efficiency. Mutation of even one of these alanine residues yielded QCascade variants with integration efficiencies 2- to 3-fold above wild-type, but interestingly, these changes did not accompany concomitant increases in PAM stringency (**Fig. 4b**). On the other hand, our episomal PAM screen in human cells revealed a wild-type ‘CN’ preference that had not previously been observed in *E. coli*, and we hypothesize that this difference may result from the larger DNA search space in the human cell milieu. The quality of our cryoEM maps also provided a detailed view of RNA-DNA base-pairing interactions, enabling visualization of energetically unfavorable nucleobase positioning along the heteroduplex (**Fig. 4d-h**). Close analysis of the surrounding Cas7 residues implicated several hydrophobic side chains in enforcing this positioning (**Supplementary Fig. 8**), and we therefore introduced mutations with less bulky side chains to potentially stabilize heteroduplex formation. Interestingly, however, most Cas7 variants complete abolished integration activity (**Fig. 4i**), suggesting that these mutations adversely affected DNA binding and/or QCascade complex formation.

Alongside our efforts at engineering specific *Pse*CAST components for DNA integration activity improvements, we considered a parallel path that would instead leverage pre-existing components from homologous CAST systems. Our previous experiments revealed the orthogonal properties of diverse type I-F CAST systems, which precluded mixing and matching of homologous components into single systems^26^, but we hypothesized that a more nuanced, structure-guided approach could reveal unique opportunities for the construction of synthetic chimeric designs that would retain key protein-protein interactions necessary for transposition. To this end, we leveraged AlphaFold^59^ to generate predicted structures of TnsA-TnsB interacting with a heptameric TnsC ring (**Fig. 5b**), and based on the resemblance to previously determined type V-K transpososome structures (**Supplementary Fig. 10a**)^39^, we envisioned that reprogramming the TnsB C-terminus could uncover functional chimeric CASTs. This hypothesis was borne out with data demonstrating that chimeric CASTs, in which the DNA targeting module of *Vch*CAST was combined with the DNA integration module of *Pse*CAST, functioned robustly for RNA-guided DNA integration (**Fig. 5**). Next, we further extended these chimeric designs to a variety of type I-F systems, and we demonstrated the first example of coordinated activity between type I-F and type V-K CAST machineries (**Supplementary Fig. 11**). Based on these promising results, we expect that future modifications will enable additional chimeric starting points for future engineering, such as at the TniQ–TnsC interface (**Supplementary Fig. 10b,c**).

The ability to coordinate targeted integration with transposase proteins derived from unique families^23^ opens the door to novel, diverse chimeric CAST designs that can sample combinatorial sequence spaces unexplored by evolution. With growing evidence that additional CAST subtypes can be leveraged for genome editing applications in human cells^61–63^, the ability to exchange modules with ease may be key for future CAST engineering efforts. Collectively, our work showcases diverse, structure-guided approaches to understand and improve CAST function, and opens the door to a far greater combinatorial space for leveraging CASTs systems as genome editing tools.

## METHODS

### Protein purification

The TniQ-Cascade complex from *Pse*CAST (*Pse*QCascade) was overexpressed and purified as previously described^20^, with the following modifications. All proteins were codon optimized and placed downstream of consensus RBS sequences, and TniQ contained an N-terminal 10xHis-TEV tag. The minimal CRISPR array was encode upstream of *cas7* and contained a 32 bp spacer targeting the AAVS1 locus (see **Supplementary Table 1** for detailed plasmid sequences). After overnight expression at 0.5 mM IPTG, cell pellets were resuspended in QCascade lysis buffer (50 mM Tris-Cl, pH 7.5, 700 mM NaCl, 0.5 mM PMSF, EDTA-free Protease Inhibitor Cocktail tablets (Roche), 1 mM dithiothreitol (DTT), 5% glycerol) and lysed by sonication. Lysates were clarified by centrifugation at 15,000 x g for 30 min at 4 °C. Initial purification was performed by immobilized metal-ion affinity chromatography with NiNTA Agarose (Qiagen) using NiNTA wash buffer (50 mM Tris-Cl, pH 7.5, 700 mM NaCl, 10 mM imidazole, 1 mM DTT, 5% glycerol) and NiNTA elution buffer (50 mM Tris-Cl pH 7.5, 700 mM NaCl, 300 mM imidazole, 1 mM DTT, 5% glycerol). The sample was further purified by size exclusion chromatography over a Superose 6 Increase 10/300 column (GE Healthcare) equilibrated with QCascade storage buffer (20 mM Tris-Cl, pH 7.5, 700 mM NaCl, 1 mM DTT, 5% glycerol). Fractions were pooled, concentrated, snap frozen in liquid nitrogen, and stored at −80 °C. TEV cleavage was not performed.

### Plasmid construction

Bacterial expression plasmids for *Pse*QCascade were codon-optimized for *E. coli* and synthesized by GenScript. For human cell transfections, genetic components encoding *Pse*CAST proteins were codon-optimized for human cells, synthesized by GenScript, and cloned into pcDNA3.1 expression vectors. All CAST constructs were cloned into plasmids using a combination of restriction digestion, ligation, Gibson assembly, and Golden Gate assembly. All PCR fragments for cloning were generated in-house using Q5 DNA Polymerase (New England Biolabs (NEB)) and gel purified using Qiagen Gel Extraction.

To clone the 4N PAM library used for HEK293T cell episomal integration assays, two overlapping oligos containing ‘NNNN’ were phosphorylated with T4 PNK (NEB) and hybridized at 95 °C for 2 min before cooling to room temperature. The resulting oligoduplex was ligated into a target plasmid vector predigested with BsmBI (55 °C for 2 h) using T4 DNA ligase (NEB). Cloning reactions were transformed into chemically competent NEB Turbo *E. coli*, plated on agar plates with the appropriate antibiotic to grow overnight, and inoculated in 5 uL LB media and antibiotic for approximately 7 h. Colony counting was then performed to ensure sufficient library diversity. Plasmids were then purified using Qiagen Miniprep columns verified by a combination of Sanger sequencing (Azenta/Genewiz) and whole-plasmid nanopore sequencing (Plasmidsaurus), and ultimately characterized by high-throughput sequencing (Illumina).

### CryoEM structure determination

Purified *Pse*QCascade was serially diluted in a modified buffer (20 mM Tris-Cl, pH 7.5, 200 mM NaCl, 1 mM DTT) for initial imaging experiments. Target DNA (NTS: 5′-TTCATCAAGCCATTGGACCGCCACAGTGGGGCCACTAGGGACAGGATTGGTGACCTTCGCCTTGACGGCCAAAA-3′, TS: 5′-TTTTGGCCGTCAAGGCGAAGCTGAAAAGCAATGAAGCCAAAGCGTCCTGTAAGGCGGTCCAATGGCTTGATGAA-3′) was duplexed by mixing the NTS and TS in equimolar concentrations, heated to 95° C, and then cooled to room temperature. 50 µM aliquots were then snap frozen. Purified *Pse*QCascade aliquots were incubated with a 5X molar excess of target DNA for 10 min at room temperature with a total reaction volume of 50 µL. The complex (2-4 µM range) was initially imaged in a Talos L120C (Thermo Fisher) electron microscope equipped with a LaB_6_ electron source and a Ceta-M camera. Negative staining experiments were carried out using uranyl-formate solution at 0.75% (w/v) in water. CF-400 (EMS) continuous carbon grids were activated for 30 s using a Ar/O_2_ gas mix plasma at 25 W using a Solarus2 plasma cleaner (Gatan). Immediately after plasma activation, 3 µL of the *Pse*QCascade/DNA complex at concentrations of 1, 2 and 4 µM were applied to the activated grids. After 1 min incubation, the excess solution was gently blotted away, and 3 µL of 0.75% uranyl-formate solution was added for an additional 1 min incubation. Excess staining solution was blotted away and the grids were left on the bench drying for 5 min. Grid screening revealed well stained, homogeneous, and dispersed particles with a circular shape compatible in dimensions and shape with the estimated molecular size of the complex, as well as showing similarities with previously reported images of other Cascade complexes (**Supplementary Fig. 2a**).

We chose the 1 µM concentration grid for manual collection of 10 negative staining images (pixel size 2.5 Å/pixel, 1 s exposure, -2 to -3 µm defocus) for exploratory class-2D analysis in Relion4. The resulting negative staining C2D averages confirmed the homogeneity of the sample and its potential for high-resolution (**Supplementary Fig. 2a, left**). Next, we explored the behavior of the complex under cryogenic conditions using the negative stain conditions as a reference starting point. We vitrified UltraAu foil 1.2/1.3 ‘Gold’ grids (Quantifoil) using a VitroBot Mark IV (Thermo Fisher) set up to 100% humidity and 4 °C. The sample concentration was in the 2–4 µM range. Grids were plasma cleaned with the same protocol described for the negative staining grids, and after application of 3 µL solution, the grids were blotted and plunged frozen in liquid ethane. Vitrobot settings were: blot force -5, drain and waiting time 0 with blotting times variating between 2.5 to 3.5 s. Following these parameters, we froze 8 grids, 4 grids at 2 µM concentration and 4 grids at 4 µM concentration. 2 grids, one at 2 µM and another at 4 µM concentration were transferred to a cooled 910 side entry holder (Gatan) for screening under cryogenic conditions in the same Talos L120C microscope used for negative staining using similar imaging conditions. Both grids showed good ice distribution, with the 2 µM grid showing better particle distribution and contrast in ice. Using SerialEM, we collected 10 images with similar settings as in negative staining experiments for exploratory reference-free C2D analysis in Relion4 under cryogenic conditions (**Supplementary Fig. 2a, middle**). The resulting C2D averages were promising, with distinctive and multiple views of the complex. The grid was recovered and stored for high resolution data collection in a Titan Krios G3i electron microscope equipped with a BioQuantum/K3 energy filter and direct detection.

High resolution data was collected at high magnification with 2x hardware binning in the K3 detector (0.6485 Å/pixel size after binning) at a fluence of ∼20e^−^/pixel/s and 1 s exposure time for a total dose of ∼50 e^−^/Å^2^. Defocus range was adjusted to vary between −0.8 to −2 µm, and the total number of K3 fractions was adjusted to 50. 24 h collection on the recovered grid yielded ∼22,000 images which were on-the-fly motion corrected in Relion4 with ctf estimation in ctffind4. Image processing was integrally done in Relion 4 and cryoDRGN. First, we manually selected 100 images for Laplacian picking, which yielded ∼4,000 particles that were normalized and extracted with 8 times binning. Fast C2D analysis using the VDAM algorithm generated C2D averages in multiple orientations that were selected and used as training set for Topaz, used through the Relion wrapper. Using the optimized trained model from Topaz, the full dataset of ∼22,000 images yielded ∼1.5 million particles that after two C2D steps using T parameters of 3 and then 6 was reduced to ∼667,000 particles. ArnA contamination accounted for the bulk of the eliminated particles. Next, we refined the reduced dataset using a filtered map of *Vch*QCascade as reference. We did not perform alignments with this initial classification (K20, tau fudge T=6).

We identified multiple classes with damaged or poorly aligned particles, a class without the TniQ dimer, and a dominating class with better features. A re-extraction step was then performed with the recenter option activated and at 4x binning (2.594 Å/pixel). After selection of 2D class averages showing secondary structure features, an ab-initio 3D model was reconstructed using the Stochastic Gradient Descent (SGD) algorithm with all selected particles from the class 2D job (K4, tau fudge T=3). A second 3D refinement produced a consensus refinement in the 5 Å range that upon inspection showed clear secondary features and substantial heterogeneity at the PAM distal region hosting the TniQ dimer. A soft-mask (10 pixel extension, 8 pixel soft edge and initial threshold of 0.002) was used for 3D classification without alignment using 20 classes and T parameters 3, 6 and 8. A minor population (∼8% of the particles) of Cascade without TniQ was identified and removed from the dataset, together with poorly aligned or damaged particles, reducing the total dataset to ∼128,000 particles. Re-refinement of this dataset after re-extraction to binning 2 (∼1.2 Å/pixel) produced a sub-3Å map, but exacerbated heterogeneity of the TniQ dimer region was evident.

Using focused classification of this region of the map produced multiple classes without clear discrete states, suggesting continuous heterogeneity. Before applying a multibody approach, we re-refined the ∼128,000 particle dataset after refining the ctf parameters (defocus values per particle and astigmatism per micrograph) followed by Bayesian particle polishing for signal decay and local particle movement correction. We defined via soft masking (6 pixel mask extension, 6 pixel soft edge decay, initial threshold 0.002) three rigid body groups: the first body included Cas8, and the first Cas7 monomer (Cas7.1), the second body contained Cas7 monomers 2 to 5, and the third body included the TniQ dimer, Cas6, Cas7.6, and the crRNA 3′-proximal hairpin. Residual rotation priors were defined to 10 degrees with translation offset of 2 pixels. We designed two wide masks: one (body 1) covering the best part of the map and including Cas8, the first five Cas7 proteins, and surrounding densities including the corresponding sections of the crRNA-DNA heteroduplex; and a second soft mask (body 2) covering Cas7.6, Cas6, and the TniQ dimer. Multibody refinement produced maps with exceptional quality for each body, with clear sub 3Å features for the Cas8 and the Cas7 regions. The maps for the PAM-distal body, including the TniQ dimer, improved substantially, but residual heterogeneity remained, especially at the distal end of the TniQ dimer.

We used ModelAngelo^64^ for initial model building using the improved maps from the multibody analysis. With default options and sequence information from the cloned constructs, ModelAngelo correctly built approximately 90% of the residues. Manual inspection of the built model corrected limited errors and completed areas where the resolution did not allow accurate placement of side chains. The built models were refined against the multibody maps independently, first with phenix refine (secondary structure restrain activated) and then with Refmac5, adjusting the experimental/ideal geometry weights manually to avoid overfitting. CryoDRGN analysis was performed with the final set of ∼128,000 particles used for multibody analysis in Relion. This set of particles was re-extracted to a box size of 128 pixels and an initial training in 1 dimension (Zdim=1) was performed. After assessing the homogeneity of this set of particles, 3 different training were performed with 2, 4 and 8 dimensions (Zdim=2, 4 and 8). Principal component analysis (PCA), UMAP, and K-means clustering dimensionality reduction techniques were used to explore the derived latent spaces, producing similar results irrespective of the Zdim used. We perform a final training with particle re-extracted to 256 pixels size and Zdim 2 and 8. Exploration of the latent space derived from these training revealed multiple conformations of the TniQ dimer, as shown in **Supplementary Figure 3**.

### Mammalian cell culture and transfections

HEK293T cells were cultured at 37 °C and 5% CO_2_ and maintained in DMEM media with 10% FBS and 100 U/mL of penicillin and streptomycin (Thermo Fisher Scientific). 24 h before transfection, a 48-well plate was coated with poly-D-lysine (Thermo Fisher Scientific) and seeded with 10,000 cells per well. Cells were transfected with DNA mixtures and 1 μL of Lipofectamine 2000 (Thermo Fisher Scientific) per the manufacturer’s instructions. Transcriptional activation and integration assays were performed as previously described^17^. For plasmid-based PAM library assays, cells were co-transfected with the following *Pse*CAST CAST plasmids: 200 ng pTnsAB, 50 ng pTnsC, 75 ng pQCascade, 100 ng pCRISPR (crRNA), 200 ng pDonor, and 100 ng pTarget (4N PAM library). Cells were harvested 4 days after transfection using previously described methods^17^.

### Analysis of HEK293T integration assays

Genomic integration assays were analyzed as previously described^17^. In brief, 5 µL of genomic lysate (10% of total lysate volume) was used for 2 rounds of PCR. In the first PCR, a forward primer was used that anneals to the AAVS1 locus, and a reverse primer was used that anneals to both the AAVS1 locus and a primer binding site in the donor DNA (see **Supplementary Table 3** for oligonucleotide sequences). These oligos included 5′ overhangs encoding read 1 and read 2 Illumina adapters. In the second PCR, ‘universal’ primers were used, which anneal to the read 1 and read 2 sequences and append unique index sequences and the remaining Illumina adapter sequences for next generation sequencing. Samples were then pooled, gel purified, and sequenced on a NextSeq 500/550 with at least 75 cycles in read 1. The relative abundance of reads that contain a *Pse*CAST transposon end sequence (representing an integration read) vs. downstream AAVS1 sequence (unintegrated read) was calculated.

For the episomal PAM library assay, samples were prepared as above except a different forward oligo was used that anneals directly upstream of the degenerate PAM library in PCR 1, such that we would capture both the PAM sequence and the presence of the transposon end sequence with the forward read (see **Supplementary Table 3** for oligonucleotide sequences). PCR 1 cycles were reduced to 15 cycles. After Illumina sequencing, reads were filtered to have a transposon end sequence, thus representing a PAM library member which was successfully targeted by *Pse*CAST for DNA integration. The input library was sequenced as well, to calculate enrichment and depletion scores. Library members were then ranked by their enrichment values (proportion of output library / proportion of input library). The top 10% of library members were used to generate a consensus WebLogo (Version 2.8.2, 2005-09-08, weblogo.berkeley.edu) for the PAM preference of each Cas8 variant. All library members and their associated enrichment values were used to generate PAM wheels using Krona^65^.

### *E. coli* repression and integration assays

*E. coli* transcriptional repression assays were performed as previously described^40,57^, with some minor modifications. In brief, an *E. coli* strain expressing mRFP from the chromosome, a gift from L. S. Qi, was transformed with pQCascade. We initially attempted to use pQCascade plasmids with a strong J23119 promoter, but due to toxicity associated with strong *Pse*QCascade expression, we switched to a weaker J23101 promoter for all pQCascade constructs. We designed crRNA sequences to target the template strand of mRFP proximal to the 5′ end of the coding region (60 bp downstream of the mRFP start codon). Two replicates were performed for each unique transformation, and relative mRFP repression was analyzed as previously described^40^.

Integration assays were performed as previously described^15,40^, with the following modifications. Although J23101 promoters were used for QCascade, J23119 promoters were still used for constitutive expression of all TnsABC cassettes, as there was no observed toxicity. In brief, TnsABC expression vectors harboring donor DNA (pDonor-TnsABC) encoded a *tnsA-tnsB-tnsC* operon downstream of a strong constitutive promoter (J23119), as well as a mini-transposon donor DNA of 0.9 and 1.2 kb in length for *Vch*CAST and *Pse*CAST, respectively, all on a pUC19 backbone. Strains harboring medium-strength J23101 promoter-controlled pQCascade constructs were first made chemically competent, followed by duplicate transformations with pDonor-TnsABC and lysate generation for qPCR after an 18 h incubation at 37 °C. Lysates were analyzed via qPCR, as previously performed^15,40^.

## Data availability

Cryo-EM maps and models will be deposited on EMDB and PDB and released upon publication. Source data for protein gels are included as **Supplementary Fig. 12**.

## Author Contributions

G.D.L., A.R.L., and S.H.S. conceived of and designed the project. G.D.L. purified *Pse*QCascade. G.D.L. and A.R.L. performed all cellular experiments and cellular experimental analyses, with the exception of *E. coli* repression and integration assays, which were performed by D.J.Z and A.R.L. I.S.F. collected cryoEM data and performed structure determination. G.D.L., A.R.L., I.S.F., and S.H.S. discussed the data and wrote the manuscript, with input from D.J.Z.

## Acknowledgements

We thank Z. Akhtar for laboratory support, R.T.K. for assistance in mammalian assay design and cloning, L. F. Landweber for qPCR instrument access, the Columbia Stem Cell Initiative Flow Cytometry Core, the JP Sulzberger Columbia Genome Center for NGS support, and the CryoEM facility of the St. Jude Children’s Research Hospital in Memphis, Tennessee, USA where the cryoEM high resolution data was collected. Some of this work was performed at the National Center for CryoEM Access and Training (NCCAT) and the Simons Electron Microscopy Center located at the New York Structural Biology Center is supported by the NIH Common Fund Transformative High Resolution Cryo-Electron Microscopy program (U24 GM129539,) and by grants from the Simons Foundation (SF349247) and NY State Assembly. S.H.S. was supported by NIH grant DP2HG011650, a Pew Biomedical Scholarship, a Sloan Research Fellowship, an Irma T. Hirschl Career Scientist Award, and a generous startup package from the Columbia University Irving Medical Center Dean’s Office and the Vagelos Precision Medicine Fund.

## SUPPLEMENTARY FIGURES

**Supplementary Figure 1.**
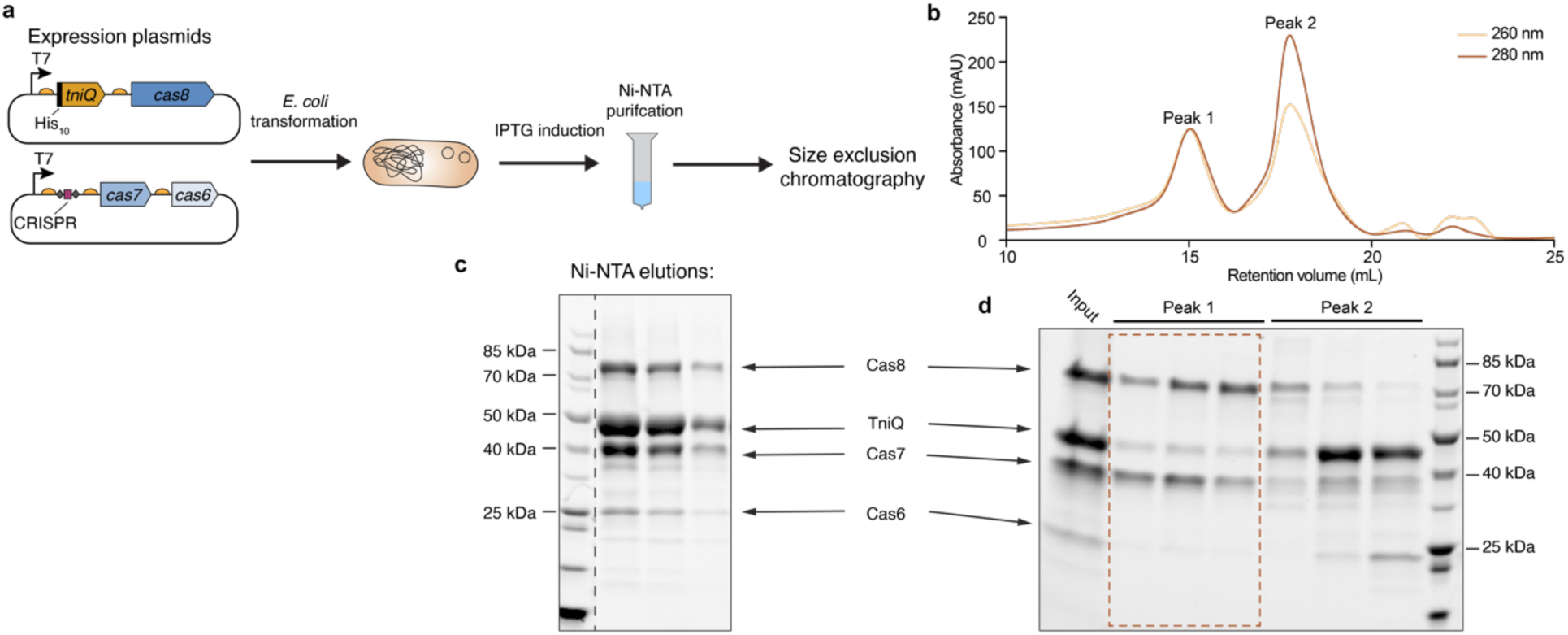
Purification of Qcascade from *Pse*CAST (Tn*7016*). **a,** The two schematized expression plasmids (left) encode *E. coli* codon-optimized *Pse*CAST QCascade genes and a crRNA cassette, with a. strong ribosome binding site (half-circle) upstream of each protein-coding gene. After transformation of BL21(DE3) cells and IPTG induction, *Pse*QCascade was purified via Ni-NTA affinity chromotography and size exclusion chromatography (SEC). Codon-optimized expression plasmids were used after the native operon failed to generate detectable QCascade complexes after SEC. **b,** SEC chromatogram of *Pse*QCascade showing 2 distinct peaks. **c,** SDS-PAGE gel of representative Ni-NTA elution fractions that were pooled and used for SEC. QCascade subunits are labeled. **d,** SDS-PAGE gel of both peaks from SEC. Elutions from peak 1, marked with a red dashed box, were pooled and used for cryoEM.

**Supplementary Figure 2.**
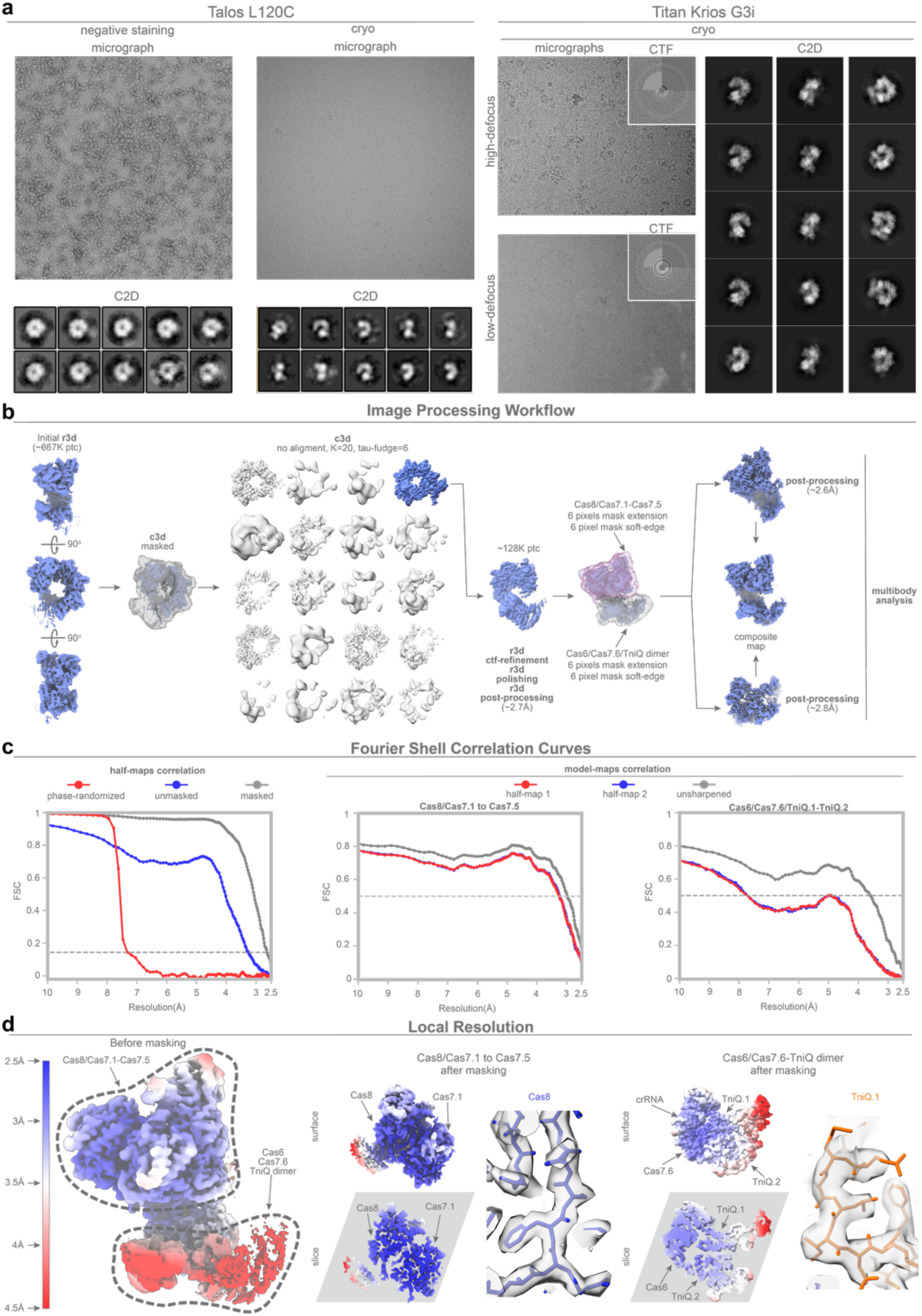
CryoEM imaging, data processing, and model refinement. **a**, Preliminary sample characterization for cryoEM grid optimization. Left, Talos L120C microscope analysis showing exemplary negative staining micrograph (left) and cryogenic micrograph (right). Corresponding reference-free 2D class averages from particles obtained from 10 images are shown below each image, with a calibrated pixel size of 2.5 Å. Right, two grids from the Talos L120C screening were recovered and loaded into a Titan Krios G3i microscope, and a large dataset was collected at a pixel size of 0.644 Å. Two images at different defoci are shown with their corresponding CTF images (inset). Reference-free 2D class averages are shown on the right, with multiple different views revealing details compatible with protein secondary structure. **b**, Image processing workflow implemented in Relion4 for high-resolution structure determination. Briefly, from left to right: an ab-initio 3D model was reconstructed after selection of 2D class averages; using this as a reference, a consensus refinement was generated; inspection of this preliminary map revealed heterogeneity, especially in the region of the TniQ dimer (**Methods**). However, after unbinning and multiple rounds of 3D refinements, the map still exhibited residual heterogeneity in the region adjacent to the Cas6 protein, suggesting mobility of the TniQ dimer with respect to Cascade. To improve the maps and to analyze TniQ dynamics, two masks were designed (**Methods**), yielding improved the densities and B-factors for the first body, but the second body exhibited a significant improvement in terms of resolution and general density quality. **c**, Fourier Shell Correlation (FSC) curves for the half-maps and model-maps. **d**, Local resolution depictions of the final map before and after the multibody approach.

**Supplementary Figure 3.**
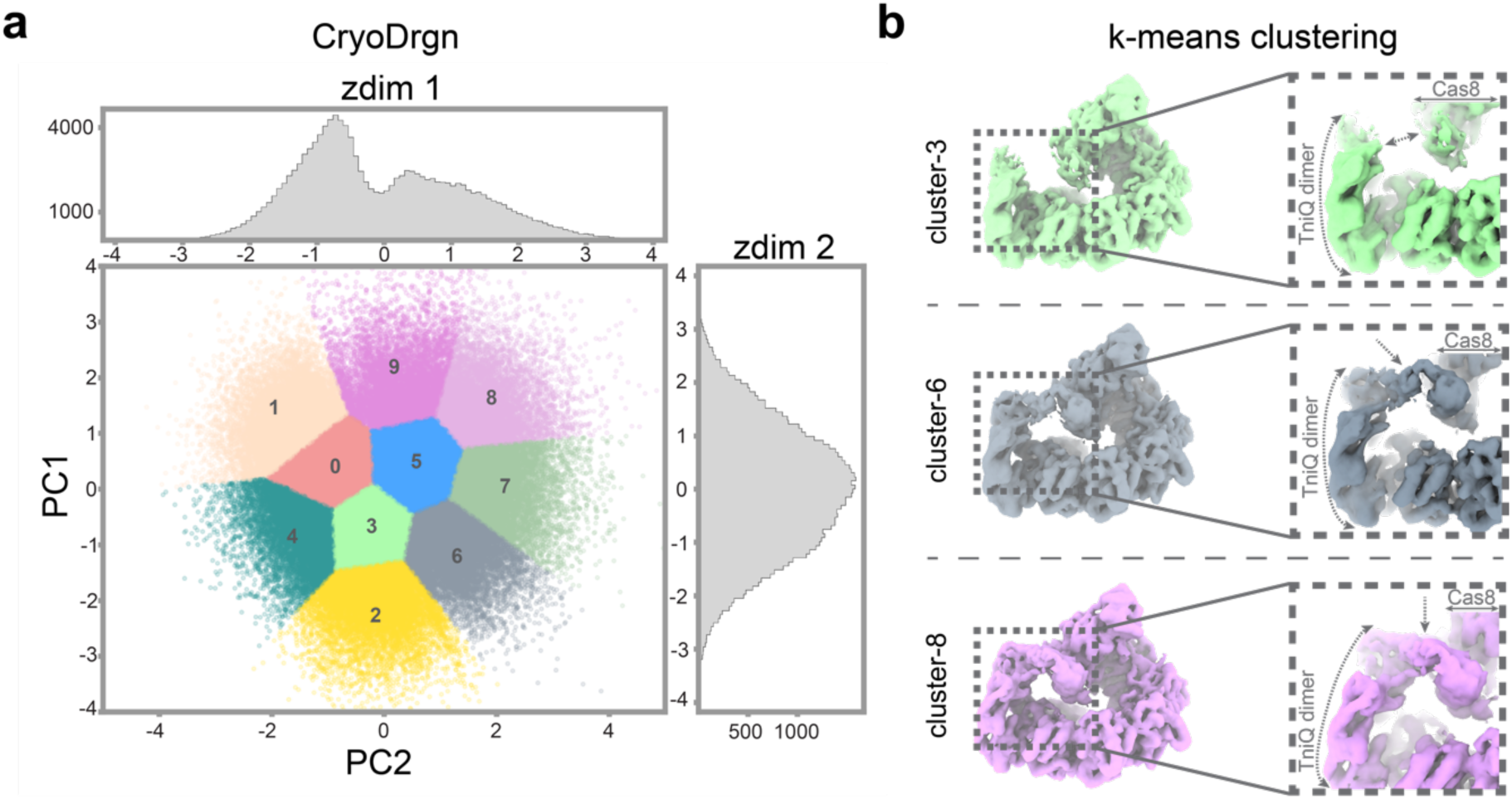
Visualization of TniQ dimer dynamics with cryoDRGN. **a,** cryoDRGN analysis, using the same set of particles (∼128,000) identified using Relion4 classifications, revealed dynamics of the TniQ dimer and uncovered multiple conformational states. We trained cryoDRGN on our dataset with multiple values of the zdim (2, 4 and 8), and found that the derived latent space for different runs was similar. Shown is a principal component analysis of the latent space derived from the run at zdim = 2. **b,** Segmentation of this latent space via k-mean clustering reveled multiple TniQ dimer conformations: an ‘open’ position, in which the TniQ dimer is distant from the Cas8 ɑ-helical domain (cluster 3, green); an intermediate position, where the distal end of the TniQ dimer marginally contacts the Cas8 ɑ-helical domain (cluster 6, grey); and a compact conformation, in which the TniQ dimer closely approaches the Cas8 ɑ-helical domain (cluster 8, pink). In all cryoDRGN-generated maps, the Cas8 ɑ-helical domain remains in a similar position and conformation. with only the TniQ dimer exhibiting pronounced fluctuations

**Supplementary Figure 4.**
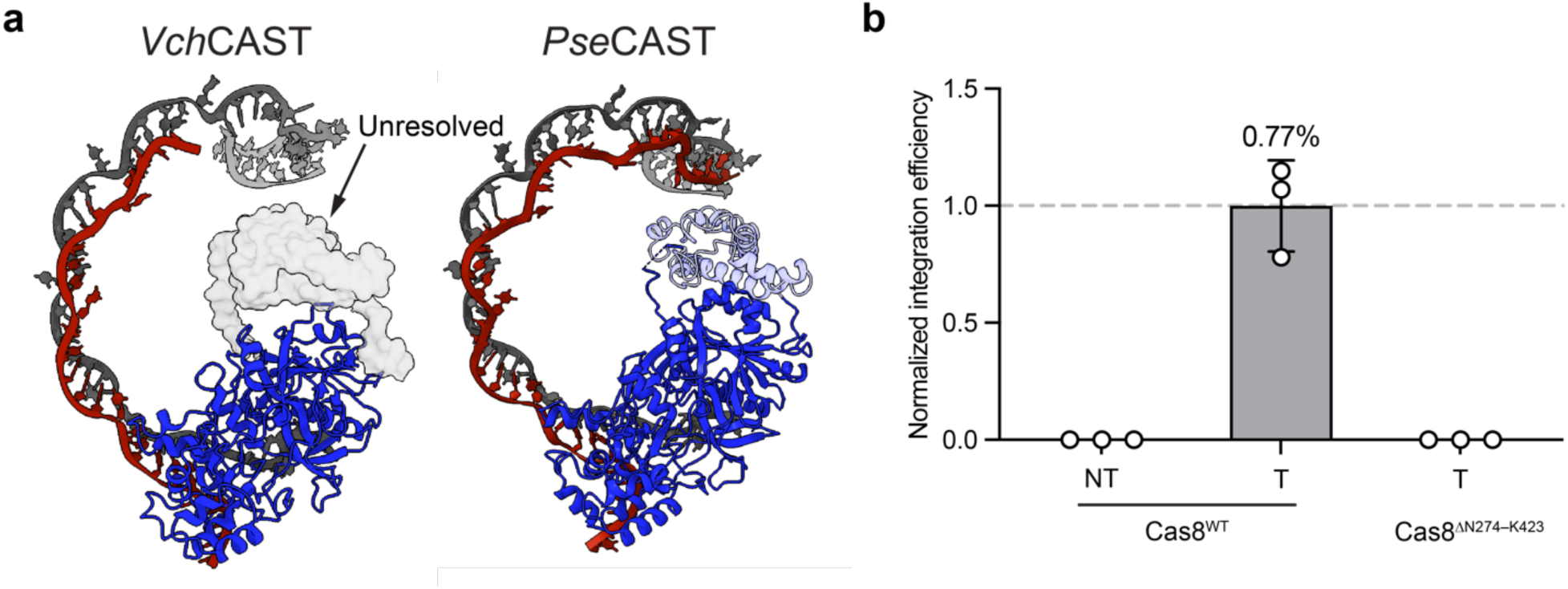
Cas8 ɑ-helical domain deletion abolishes RNA-guided DNA integration. **a**, Comparison of select regions of the DNA-bound QCascade complex from *Vch*CAST (left, PDB: 6PIJ) and *Pse*CAST (right), including the crRNA (grey), target DNA (red), and Cas8 (blue). The Cas8 ɑ-helical domain from *Pse*CAST (residues 274–423) is shown in light blue, and was replaced with a flexible, 10-amino acid GS linker in subsequent integration assays. **b**, Normalized efficiency of RNA-guided DNA integration at *AAVS1*, tested in HEK293T cells and measured by amplicon sequencing (**Methods**). Experiments used WT Cas8 and either a non-targeting (NT) or targeting (T) crRNA, or a targeting crRNA and Cas8 mutant, in which residues N274–K243 were replaced with a 10-amino acid GS linker. Data are shown as mean ± s.d. for n=3 independent biological samples.

**Supplementary Figure 5.**
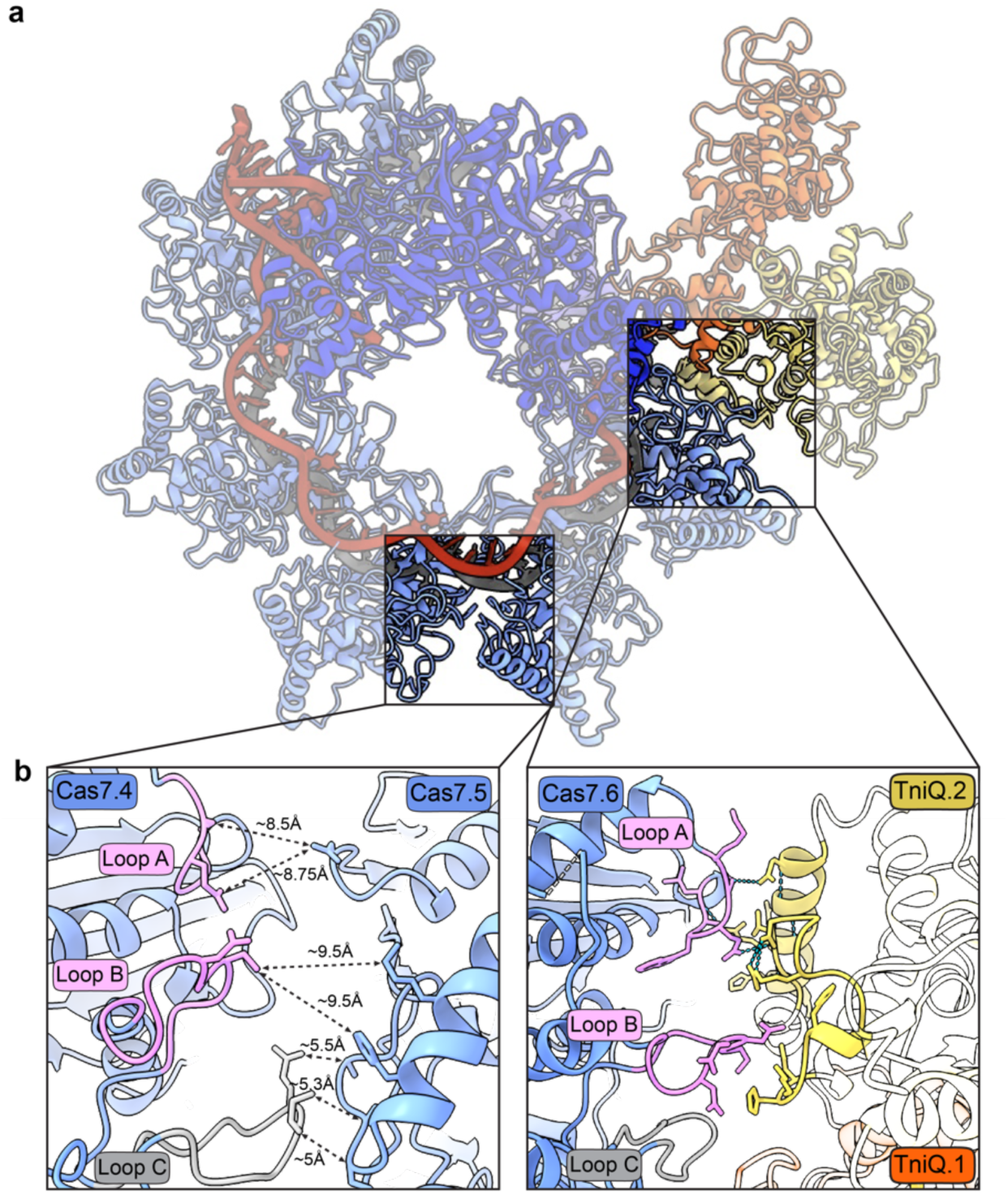
Cas7 loops chosen to selectively perturb Cas7-TniQ.2 interactions. **a,** View of overall *Pse*CAST QCascade complex, with specific regions for panel **b** highlighted. **b,** Magnified view of the different Cas7 loop interactions. Loop C participates in interactions at the interface between Cas7 monomers (left) and was therefore left intact. Amino acid sidechains in loops A and B (pink) that interact more closely with TniQ.2 were selected for mutagenesis, as detailed in **Figure 3e,f**.

**Supplementary Figure 6.**
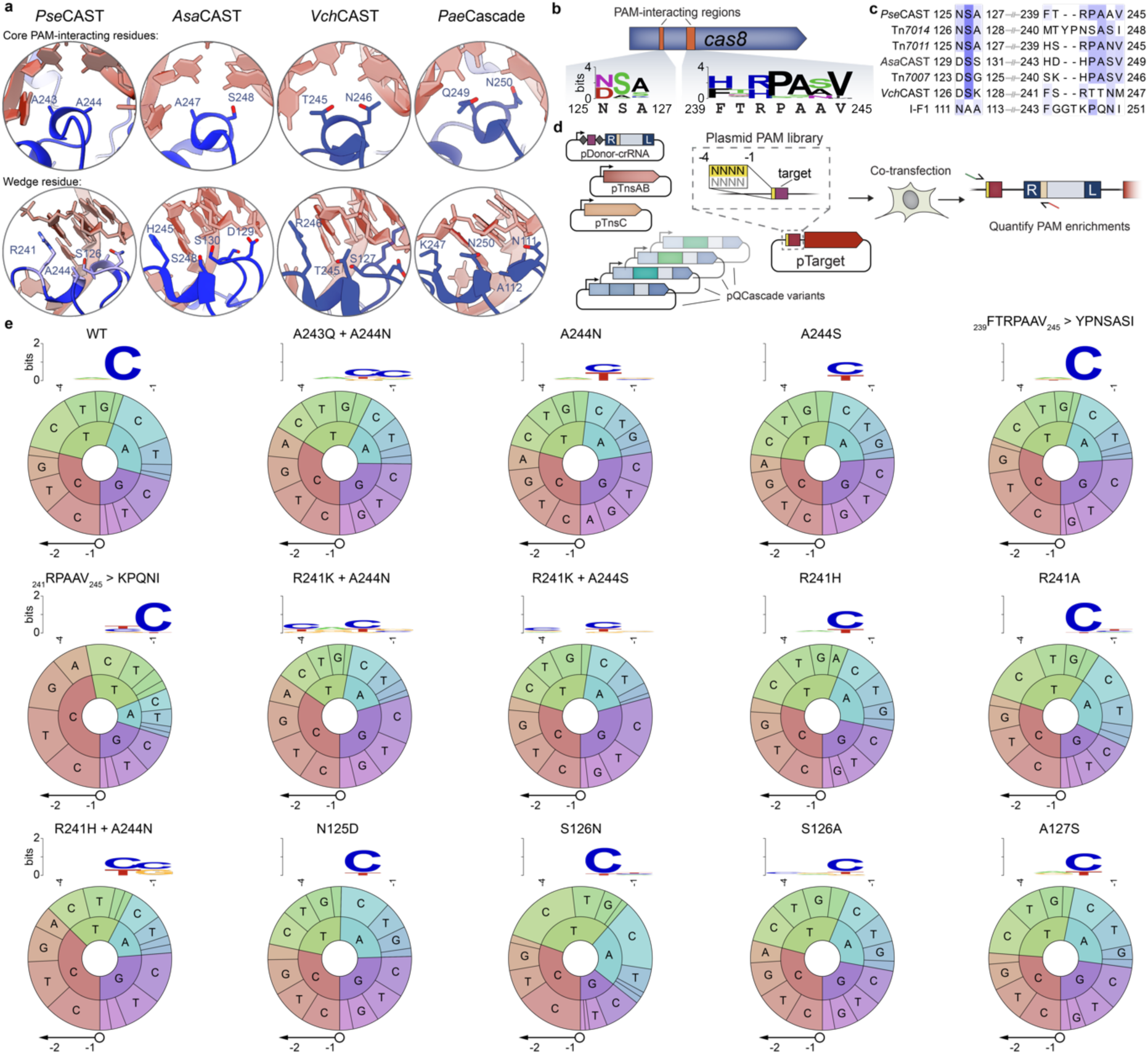
Experimental design and results for PAM library screening with Cas8. **a**, Visualization of PAM binding pockets for diverse type I-F Cascade complexes (from left to right): *Pse*CAST (this study), *Asa*CAST (PDB: 7U5D), *Vch*CAST (PDB: 6PIJ), and *Pae*Cascade (PDB: 6NE0)^47^. The top inset shows core PAM-interacting residues; the bottom inset shows the wedge residue and additional interacting residues. **b**, Amino acid sequence conservation within PAM-interacting regions of *Pse*Cas8, with the WebLogo derived from a multiple sequence alignment (MSA) of 66 homologs; the *Pae*Cas8 WT sequence is shown below. **c**, MSA of the same regions from **b**, shown for diverse type I-F Cas8 homologs from both CAST and canonical type I-F1 CRISPR-Cas systems. Conserved residues are colored in blue. **d**, Mammalian PAM library assay workflow. A target plasmid (pTarget) was generated that contains an *AAVS1* target flanked by a 4-bp randomized PAM library. Individual Cas8 mutants were screened in each transfection via a plasmid-based integration assay, in which junction PCR and next-generation sequencing revealed PAM sequences enriched within integration products (**Methods**). **e**, Detailed PAM library data for all active Cas8 variants, showing the identity of the mutation(s) (top), WebLogo of the top 10% of enriched library members (middle), and PAM wheel^65^ of all library members (bottom)^65^. The PAM wheel is displayed with the inner and outer rings representing the -1 and -2 PAM positions, respectively.

**Supplementary Figure 7.**
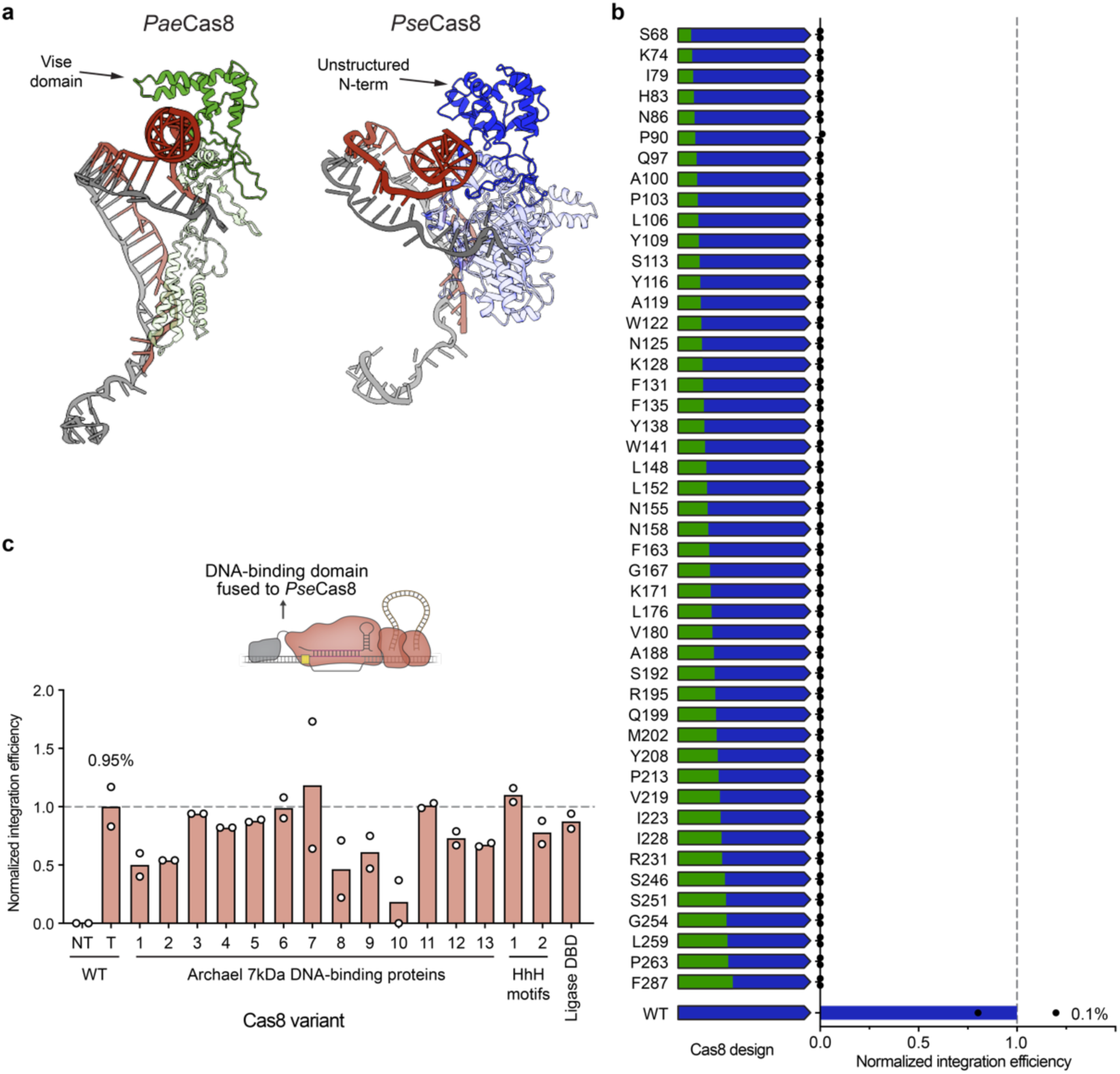
Investigating integration activity via engineering DNA-binding ability of Cas8. **a**, Comparing N-terminal regions of *Pae*Cas8 (PDB: 6NE0) and *Pse*Cas8. While *Pae*Cas8 (left) shows a vise domain clamped around the dsDNA backbone, *Pse*Cas8 shows an unstructured region at the N-terminus that does not exhibit clear dsDNA backbone interactions. **b**, Normalized RNA-guided DNA integration efficiency at *AAVS1* in HEK293T cells as measured by amplicon sequencing, for a panel of chimeric Cas8 designs in which the N-terminus of *Pae*Cas8 (type I-F1 CRISPR-Cas system, green) was grafted onto the N-terminus of *Pse*Cas8 (type I-F3 *Pse*CAST, blue); the amino acid residue listed at left indicates the graft point (*Pse*Cas8 numbering). All chimeric designs tested were non-functional for DNA integration. **c**, Normalized RNA-guided DNA integration efficiency at *AAVS1* in HEK293T cells as measured by amplicon sequencing, for a panel of Cas8 fusions designed to improve DNA binding affinity. Thirteen unique archael 7 kDa DNA-binding proteins^66^, two helix–hairpin–helix DNA binding motifs (‘HhH’)^67^, and one binding domain from *Pyrococcus abyssi* DNA ligase^52^ (‘Ligase DBD’) were tested as N-terminal *Pse*Cas8 fusions, compared to non-targeting (NT) and targeting (T) controls with WT Cas8. Data in **b** and **c** are shown as mean for n=2 biologically independent samples.

**Supplementary Figure 8.**
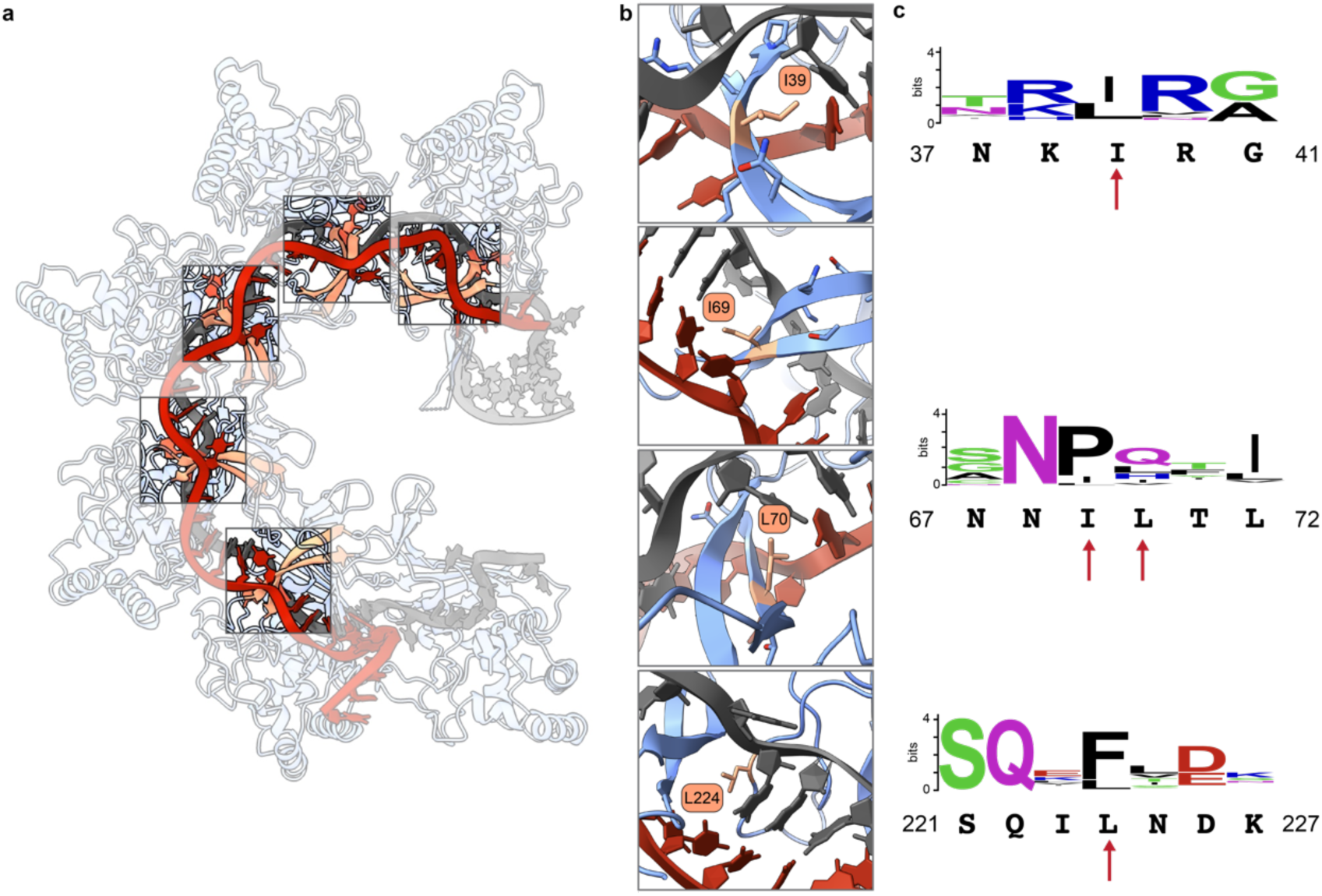
Detailed view of Cas7 interactions with the RNA-DNA heteroduplex. **a**, View of overall *Pse*QCascade complex, with the five similar Cas7-crRNA interactions highlighted. **b**, Visualization of Cas7 residues that interact with the crRNA at each flipped out nucleobase; residues with bulky and hydrophobic sidechains are highlighted and labeled. **c**, *Pse*Cas7 sequence conservation at residues in panel **b**, from a multiple sequence alignment of 98 homologs; the WT sequence is shown below the x-axis. Specific residues selected for functional investigation are marked with red arrows.

**Supplementary Figure 9.**
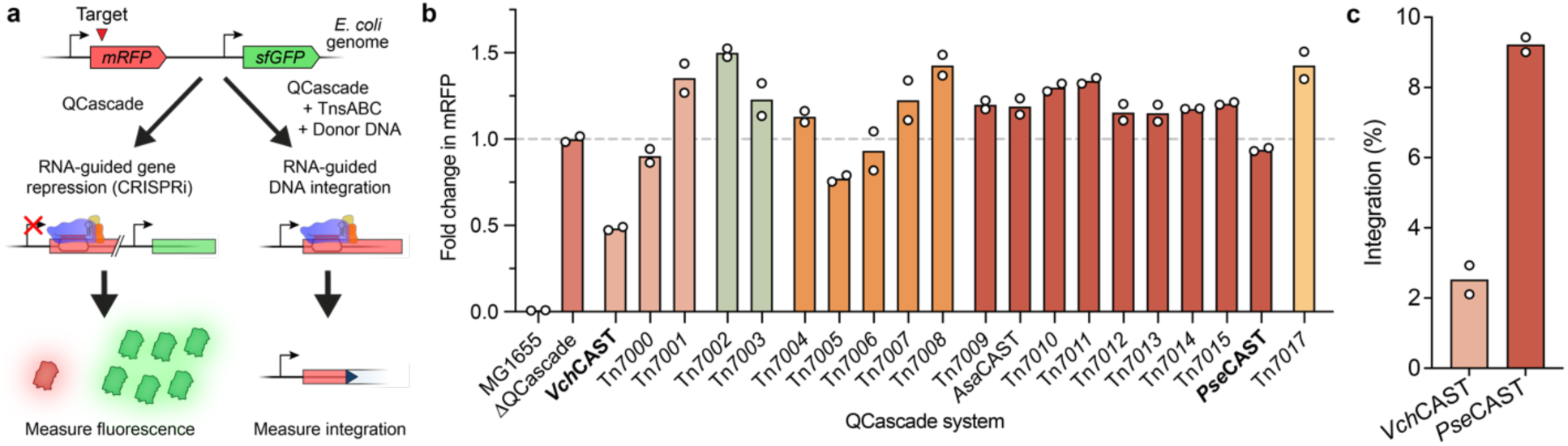
DNA binding and integration activity of diverse CAST systems in *E. coli*. **a,** Schematic of *E. coli* transcriptional repression and DNA integration assays to investigate CAST-encoded QCascade activity in bacteria. Using an engineered *E. coli* strain that constitutively expresses mRFP and sfGFP^57^, transformation of^57^. a QCascade expression plasmid driven by a medium-strength J23101 promoter leads to target DNA binding (red triangle) and mRFP repression. Alternatively, when cells are co-transformed with QCascade, TnsABC, and pDonor, RNA-guided DNA integration occurs at the mRFP target site. **b,** Bar graph showing the fold change in mRFP fluorescence for each CAST-encoded QCascade system, relative to a control experiment lacking QCascade (ΔQCascade); *Vch*CAST and *Pse*CAST are highlighted in bold text. CAST systems are colored by phylogenetic clade, as shown in **Fig. 1a. c**, Bar graph comparing DNA integration activity for *Vch*CAST and *Pse*CAST at the same mRFP target site used for repression assays, as measured by qPCR. As observed in human cells, *Pse*CAST yields higher levels of DNA integration activity despite exhibiting apparent weaker QCascade-based DNA targeting and repression. Data in **b,c** are shown as mean for n=2 independent biological samples.

**Supplementary Figure 10.**
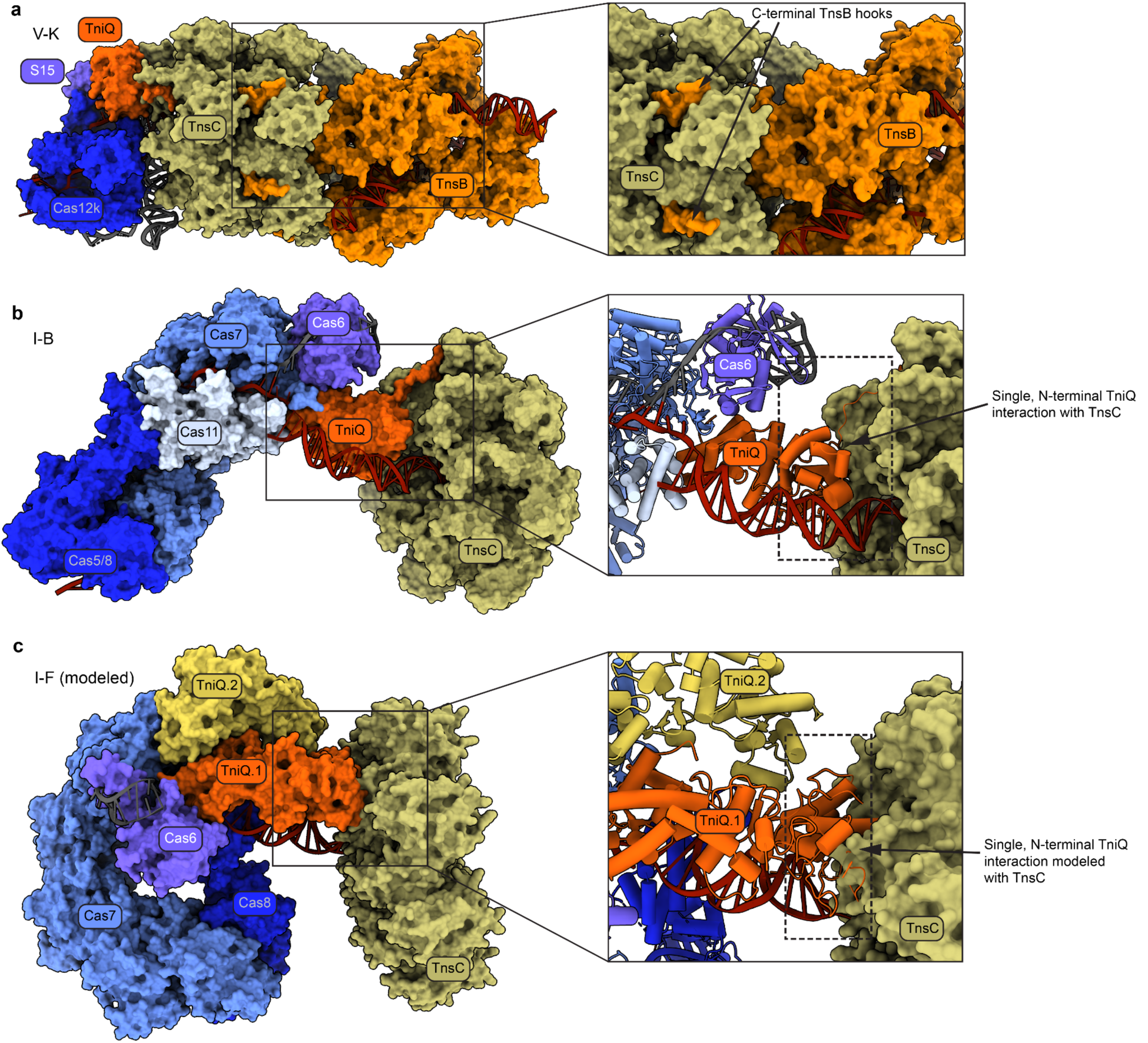
Structural inspiration for the rational design of chimeric CAST systems. **a,** The holo transpososome structure from type V-K ShCAST system (PDB 8EA3), with a magnified view (right) showing how the C-terminal hook of TnsB docks into the TnsC ATPase. **b,** The QCascade-TnsC structure from type I-B *Pmc*CAST system (PDB 8FF4), with a magnified view (right) showing the N-terminus of a monomeric TniQ interacting with the TnsC ATPase. **c,** Predicted QCascade-TnsC structure from type I-F CAST, based on previous modelling^40^ but with PDB ID: 7U5D, for which the PAM-distal DNA is better resolved. The magnified view (right) highlights the putative TniQ-TnsC interface, with the N-terminus of just one TniQ monomer within the dimeric arrangement interacting with the TnsC ATPase.

**Supplementary Figure 11.**
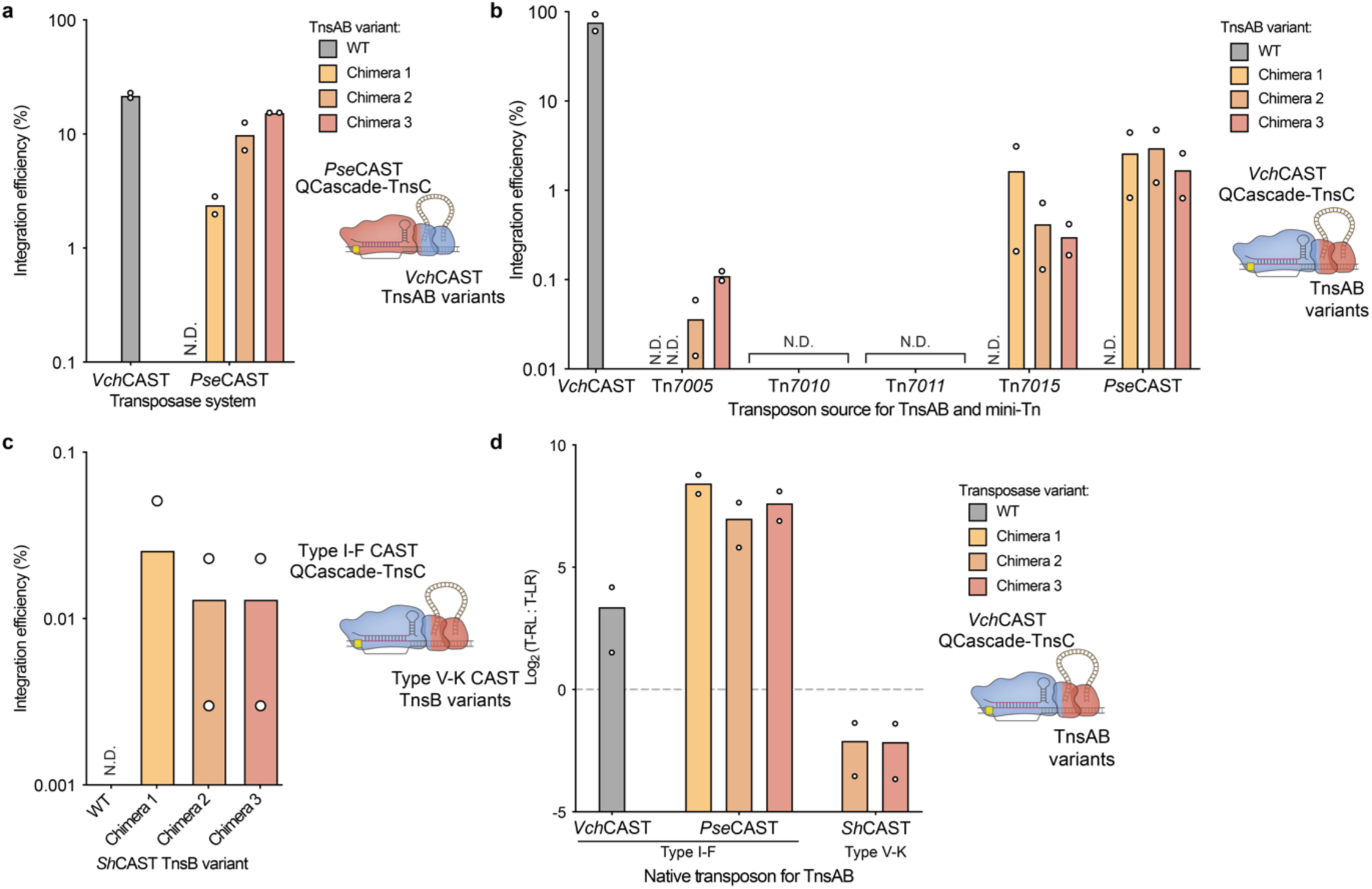
Additional chimeric TnsB designs are functional for RNA-guided DNA integration. **a**, Investigating reciprocal chimeric designs to coordinate transposition between *Pse*QCascade-TnsC and *Vch*TnsAB. WT TnsAB sequences for both *Vch*CAST and *Pse*CAST and three unique chimeric inspired by the most active variants in **Fig. 5e** (variants V585, S589, and Q594) were tested. Only chimeric TnsAB variants enabled coordinated DNA integration activity when combining *Pse*QCascade and *Pse*TnsC with *Vch*TnsAB and *Vch* mini-transposons. **b**, Exploring chimeric CASTs across multiple type I-F systems. Chimeric TnsAB variants enable coordinated transposition when combining *Vch*QCascade-TnsC with TnsAB constructs sourced from diverse Type I-F CASTs; Tn numbers were defined previously^26^. **c**, Designing chimeric CASTs across evolutionarily distinct CAST families. Chimeric *Sh*CAST TnsB constructs (inspired by functional chimeric *Pse*TnsABs) can coordinate low levels of transposition between type I-F and type V-K CAST systems. For chimera 1, only one of two biological replicates exhibited detectable integration. **d**, Insertion site orientation preference of *Vch*CAST, *Pse*CAST TnsAB chimeras, and *Sh*CAST TnsB chimeras. *Vch*CAST TnsAB and *Pse*CAST TnsAB chimeras adopt the common T-RL preference; *Sh*CAST TnsB chimeras invert the insertion site orientation preference, adopting the previously observed T-LR preference for *Sh*CAST systems^16^. Data shown as mean for n=2 independent biological samples. Chimeras 1, 2, and 3 for all homologs are listed in **Supplementary Table S2.**

**Supplementary Figure 12.**
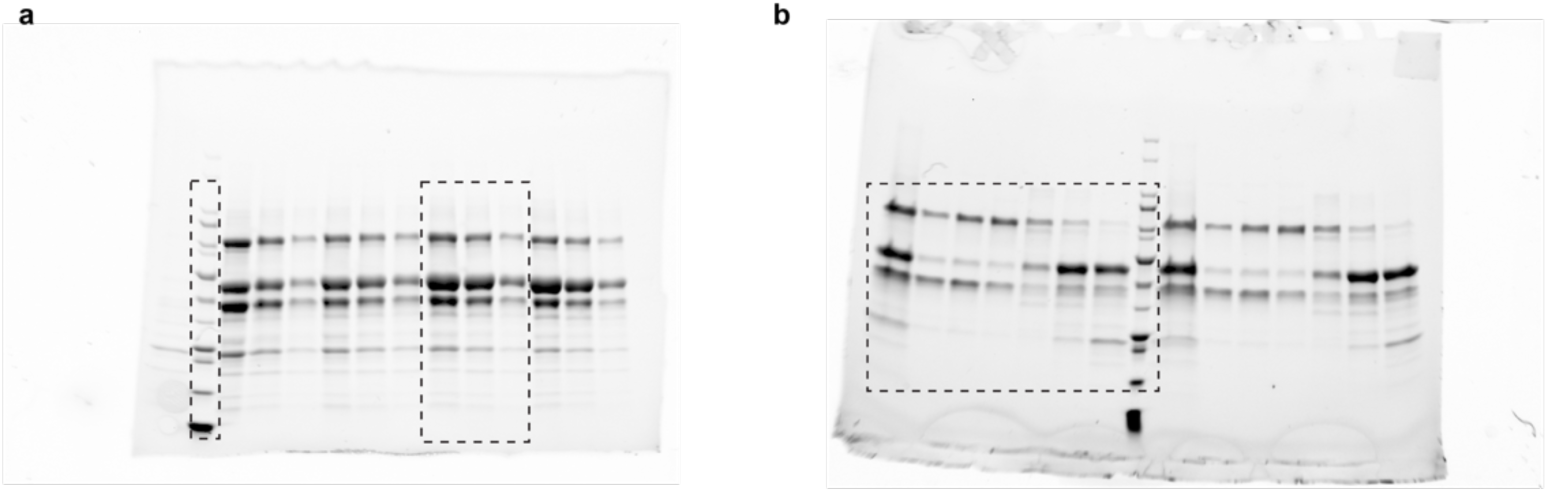
Uncropped protein gels. **a,** Uncropped image used for **Supplementary Figure 1c**. The regions shown are marked with dashed boxes. **b,** Uncropped image used for **Supplementary Figure 1d**. The regions shown are marked with a dashed box.

Supplementary Movie 1 | Conformational transitions revealed by Relion Multibody analysis defining two bodies. The first body included Cas8, the DNA-RNA duplex, and all Cas7 monomers; the second body included the TniQ dimer, Cas6, and the corresponding fragment of the crRNA interacting with Cas6.

Supplementary Movie 2 | CryoDRGN analysis of *Pse*QCascade flexibility visualized through k-means clustering of the latent space. Morphing between the two most populated states after segmentations into 20 clusters is shown.

## SUPPLEMENTARY TABLES

**Table S1. Description and sequence of plasmids used in this study.**

**Table S2. Sequence of chimeric TnsAB protein sequences used in this study.**

**Table S3: Oligonucleotides used for amplicon sequencing in this study.**

## Notes

### Competing Interest Statement

The authors have declared no competing interest.

## REFERENCES

1. Branzei, D. & Foiani, M. Regulation of DNA repair throughout the cell cycle. Nat. Rev. Mol. Cell Biol. 9, 297–308 (2008).

2. Heyer, W.-D., Ehmsen, K. T. & Liu, J. Regulation of Homologous Recombination in Eukaryotes. Annu. Rev. Genet. 44, 113–139 (2010).

3. Pawelczak, K. S., Gavande, N. S., VanderVere-Carozza, P. S. & Turchi, J. J. Modulating DNA Repair Pathways to Improve Precision Genome Engineering. ACS Chem. Biol. 13, 389–396 (2018).

4. Kanca, O. et al. An efficient CRISPR-based strategy to insert small and large fragments of DNA using short homology arms. eLife 8, e51539 (2019).

5. Zuccaro, M. V. et al. Allele-Specific Chromosome Removal after Cas9 Cleavage in Human Embryos. Cell 183, 1–15 (2020).

6. Adikusuma, F. et al. Large deletions induced by Cas9 cleavage. Nature 560, E8–E9 (2018).

7. Kosicki, M., Tomberg, K. & Bradley, A. Repair of double-strand breaks induced by CRISPR– Cas9 leads to large deletions and complex rearrangements. Nat. Biotechnol. 36, (2018).

8. Leibowitz, M. L. et al. Chromothripsis as an on-target consequence of CRISPR–Cas9 genome editing. Nat. Genet. 53, 895–905 (2021).

9. Nahmad, A. D. et al. Frequent aneuploidy in primary human T cells after CRISPR–Cas9 cleavage. Nat. Biotechnol. 40, 1807–1813 (2022).

10. Tsuchida, C. A. et al. Mitigation of chromosome loss in clinical CRISPR-Cas9-engineered T cells. Cell 186, 4567–4582.e20 (2023).

11. Komor, A. C., Kim, Y. B., Packer, M. S., Zuris, J. A. & Liu, D. R. Programmable editing of a target base in genomic DNA without double-stranded DNA cleavage. Nature 533, 420–424 (2016).

12. Kim, Y. B. et al. Increasing the genome-targeting scope and precision of base editing with engineered Cas9-cytidine deaminase fusions. Nat. Biotechnol. 35, 371–376 (2017).

13. Gaudelli, N. M. et al. Programmable base editing of T to G C in genomic DNA without DNA cleavage. Nature 551, 464–471 (2017).

14. Anzalone, A. V. et al. Search-and-replace genome editing without double-strand breaks or donor DNA. Nature 576, 149–157 (2019).

15. Klompe, S. E., Vo, P. L. H., Halpin-Healy, T. S. & Sternberg, S. H. Transposon-encoded CRISPR–Cas systems direct RNA-guided DNA integration. Nature 571, 219–225 (2019).

16. Strecker, J. et al. RNA-guided DNA insertion with CRISPR-associated transposases. Science 364, 48–53 (2019).

17. Lampe, G. D. et al. Targeted DNA integration in human cells without double-strand breaks using CRISPR-associated transposases. Nat. Biotechnol. 42, 87–98 (2023).

18. Saito, M. et al. Dual modes of CRISPR-associated transposon homing. Cell 184, 2441–2453.e18 (2021).

19. Hsieh, S. & Peters, J. E. Discovery and characterization of novel type I-D CRISPR-guided transposons identified among diverse Tn7-like elements in cyanobacteria. 51, 765–782 (2023).

20. Halpin-Healy, T. S., Klompe, S. E., Sternberg, S. H. & Fernández, I. S. Structural basis of DNA targeting by a transposon-encoded CRISPR–Cas system. Nature 577, 271–274 (2020).

21. Wang, S., Gabel, C., Siddique, R., Klose, T. & Chang, L. Molecular mechanism for Tn7-like transposon recruitment by a type I-B CRISPR effector. Cell 186, 4204–4215.e19 (2023).

22. Park, J. U. et al. Multiple adaptations underly co-option of a CRISPR surveillance complex for RNA-guided DNA transposition. Mol. Cell 83, 1827–1838.e6 (2023).

23. Faure, G. et al. CRISPR–Cas in mobile genetic elements: counter-defence and beyond. Nat. Rev. Microbiol. 17, 513–525 (2019).

24. Vo, P. L. H., Acree, C., Smith, M. L. & Sternberg, S. H. Unbiased profiling of CRISPR RNA-guided transposition products by long-read sequencing. Mob. DNA 12, 1–17 (2021).

25. Schmitz, M., Querques, I., Oberli, S., Chanez, C. & Jinek, M. Structural basis for the assembly of the type V CRISPR-associated transposon complex. Cell 185, 4999–5010.e17 (2022).

26. Klompe, S. E. et al. Evolutionary and mechanistic diversity of Type I-F CRISPR-associated transposons. Mol. Cell 82, 616–628.e5 (2022).

27. Roberts, A., Nethery, M. A. & Barrangou, R. Functional characterization of diverse type I-F CRISPR-associated transposons. Nucleic Acids Res. 50, 11670–11681 (2022).

28. Rybarski, J. R., Hu, K., Hill, A. M., Wilke, C. O. & Finkelstein, I. J. Metagenomic discovery of CRISPR-associated transposons. Proc. Natl. Acad. Sci. U. S. A. 118, (2021).

29. Petassi, M. T., Hsieh, S. & Peters, J. E. Guide RNA Categorization Enables Target Site Choice in Tn7-CRISPR-Cas Transposons. Cell 183, 1757–1771.e18 (2020).

30. Walker, M. W. G., Klompe, S. E., Zhang, D. J. & Sternberg, S. H. Novel molecular requirements for CRISPR RNA-guided transposition. Nucleic Acids Res. 51, 4519–4535 (2023).

31. Vo, P. L. H. et al. CRISPR RNA-guided integrases for high-efficiency, multiplexed bacterial genome engineering. Nat. Biotechnol. 39, 480–489 (2021).

32. Rubin, B. E. et al. Species- and site-specific genome editing in complex bacterial communities. Nat. Microbiol. 7, 34–47 (2022).

33. George, J. T. et al. Mechanism of target site selection by type V-K CRISPR-associated transposases. Science 382, (2023).

34. Strecker, J., Ladha, A., Makarova, K. S., Koonin, E. V. & Zhang, F. Response to Comment on “RNA-guided DNA insertion with CRISPR-associated transposases”. Science 368, 1–2 (2020).

35. Tou, C. J., Orr, B. & Kleinstiver, B. P. Precise cut-and-paste DNA insertion using engineered type V-K CRISPR-associated transposases. Nat. Biotechnol. (2023) doi:10.1038/s41587-022-01574-x.

36. Park, J. U. et al. Structural basis for target site selection in RNA-guided DNA transposition systems. Science 373, 768–774 (2021).

37. Park, J. U., Tsai, A. W. L., Chen, T. H., Peters, J. E. & Kellogg, E. H. Mechanistic details of CRISPR-associated transposon recruitment and integration revealed by cryo-EM. Proc. Natl. Acad. Sci. U. S. A. 119, 1–9 (2022).

38. Querques, I., Schmitz, M., Oberli, S., Chanez, C. & Jinek, M. Target site selection and remodelling by type V CRISPR-transposon systems. Nature (2021) doi:10.1038/s41586-021-04030-z.

39. Park, J. U. et al. Structures of the holo CRISPR RNA-guided transposon integration complex. Nature 613, 775–782 (2023).

40. Hoffmann, F. T. et al. Selective TnsC recruitment enhances the fidelity of RNA-guided transposition. Nature 609, 384–393 (2022).

41. Jia, N., Xie, W., de la Cruz, M. J., Eng, E. T. & Patel, D. J. Structure–function insights into the initial step of DNA integration by a CRISPR–Cas–Transposon complex. Cell Res. 30, 182–184 (2020).

42. Wang, B., Xu, W. & Yang, H. Structural basis of a Tn7-like transposase recruitment and DNA loading to CRISPR-Cas surveillance complex. Cell Res. 30, 185–187 (2020).

43. Li, Z., Zhang, H., Xiao, R. & Chang, L. Cryo-EM structure of a type I-F CRISPR RNA guided surveillance complex bound to transposition protein TniQ. Cell Res. 30, 179–181 (2020).

44. Zhong, E. D., Bepler, T., Berger, B. & Davis, J. H. CryoDRGN: reconstruction of heterogeneous cryo-EM structures using neural networks. Nat. Methods 18, 176–185 (2021).

45. Moreb, E. A., Hutmacher, M. & Lynch, M. D. CRISPR-Cas ‘non-Target’ Sites Inhibit On-Target Cutting Rates. CRISPR J. 3, 550–561 (2020).

46. Tuminauskaite, D. et al. DNA interference is controlled by R-loop length in a type I-F1 CRISPR-Cas system. BMC Biol. 18, 1–16 (2020).

47. Rollins, M. C. F. et al. Structure Reveals a Mechanism of CRISPR-RNA-Guided Nuclease Recruitment and Anti-CRISPR Viral Mimicry. Mol. Cell 74, 132–142.e5 (2019).

48. Guo, T. W. et al. Cryo-EM Structures Reveal Mechanism and Inhibition of DNA Targeting by a CRISPR-Cas Surveillance Complex. Cell 171, 414–426.e12 (2017).

49. Chowdhury, S. et al. Structure Reveals Mechanisms of Viral Suppressors that Intercept a CRISPR RNA-Guided Surveillance Complex. Cell 169, 47–57.e11 (2017).

50. Wang, Y. et al. A novel strategy to engineer DNA polymerases for enhanced processivity and improved performance in vitro. Nucleic Acids Res. 32, 1197–1207 (2004).

51. de Vega, M., Lázaro, J. M., Mencía, M., Blanco, L. & Salas, M. Improvement of φ29 DNA polymerase amplification performance by fusion of DNA binding motifs. Proc. Natl. Acad. Sci. U. S. A. 107, 16506–16511 (2010).

52. Oscorbin, I. P., Wong, P. F., Boyarskikh, U. A., Khrapov, E. A. & Filipenko, M. L. The attachment of a DNA-binding Sso7d-like protein improves processivity and resistance to inhibitors of M-MuLV reverse transcriptase. FEBS Lett. 594, 4338–4356 (2020).

53. Tong, C. L., Kanwar, N., Morrone, D. J. & Seelig, B. Nature-inspired engineering of an artificial ligase enzyme by domain fusion. Nucleic Acids Res. 50, 11175–11185 (2022).

54. Jackson, R. N. et al. Crystal structure of the CRISPR RNA–guided surveillance complex from Escherichia coli. Science 345, 1473–1479 (2014).

55. Xue, C., Zhu, Y., Zhang, X., Shin, Y. K. & Sashital, D. G. Real-Time Observation of Target Search by the CRISPR Surveillance Complex Cascade. Cell Rep. 21, 3717–3727 (2017).

56. Aldag, P. et al. Dynamic interplay between target search and recognition for a Type I CRISPR-Cas system. Nat. Commun. 14, (2023).

57. Qi, L. S. et al. Repurposing CRISPR as an RNA-γuided platform for sequence-specific control of gene expression. Cell 152, 1173–1183 (2013).

58. Hoffmann, F. T. et al. Selective recruitment of the AAA + ATPase TnsC increases the fidelity of Type I-F CRISPR RNA-guided transposition. Manuscr. Revis. 1–60 (2021).

59. Jumper, J. et al. Highly accurate protein structure prediction with AlphaFold. Nature 596, 583–589 (2021).

60. Skelding, Z., Sarnovsky, R. & Craig, N. L. Formation of a nucleoprotein complex containing Tn7 and its target DNA regulates transposition initiation. EMBO J. 21, 3494–3504 (2002).

61. Zhang, F., Saito, M. & Faure, G. Type I-B CRISPR-Associated Transposase Systems. (2024).

62. Metagenomi Technologies. S-1. (2024).

63. Strecker, J., Zhang, F. & Ladha, A. CRISPR-associated transposase systems and methods of use thereof.

64. Jamali, K. et al. Automated model building and protein identification in cryo-EM maps. Nature (2024) doi:10.1038/s41586-024-07215-4.

65. Ondov, B. D., Bergman, N. H. & Phillippy, A. M. Krona-385.pdf. BMC Bioinformatics 385, (2011).

66. Kalichuk, V. et al. The archaeal “7 kDa DNA-binding” proteins: extended characterization of an old gifted family. Sci. Rep. 6, 37274 (2016).

67. Shao, X. Common fold in helix-hairpin-helix proteins. Nucleic Acids Res. 28, 2643–2650 (2000).

